# Shared QTLs underlie loss of a conserved trait in replicate mapping populations of *Arabidopsis thaliana*

**DOI:** 10.64898/2026.07.14.738505

**Authors:** Madison Plunkert, Ousseini Issaka Salia, Esther Woolcock, Samuel Pérez, Jeffrey Conner

## Abstract

The evolutionary mechanisms leading to nonfunctional trait loss are not well understood. Some *Arabidopsis thaliana* populations display incomplete loss of short stamens, floral organs that do not contribute significantly to seed set in this species. In nature, short stamen number is correlated with ovule number and flowering time, suggesting that tight linkage or pleiotropy with these traits could drive stamen number evolution. To investigate the role of colocalized QTLs in short stamen number evolution and the repeatability of QTL co-localization between replicate mapping populations, we performed QTL mapping of stamen number, ovule number, and flowering time for two sets of recombinant inbred lines (RILs), Belm-12 x Roda-47 and Tsu-1 x Kas-1. All three traits were correlated in Belm-12 x Roda-47 RILs, but none were correlated in Tsu-1 x Kas-1 RILs. For both stamen and ovule number, some QTLs were unique to each RIL set and others were shared. Notably, for Belm-12 x Roda-47 RILs, we mapped QTLs affecting ovule and short stamen number to a region of chromosome 5 that also affects flowering time. For Tsu-1 x Kas-1 RILs, we identified the same chromosome 5 region as a short stamen number QTL, but ovule number and flowering time QTLs map to other genomic regions. Tests for pleiotropy revealed that the flowering time and short stamen number QTLs on chromosome 5 in Belm-12 x Roda-47 are likely separate QTLs, linked but not pleiotropic, but each may have pleiotropic effects on ovule number. Thus, the large-effect stamen number QTL on chromosome 5 was consistent between crosses, but only in Belm-12 x Roda-47 does this QTL co-localize with the other two traits.

**Article summary:** Nonfunctional trait loss is common over evolutionary time. We investigated the evolutionary mechanisms influencing nonfunctional trait loss by examining the genetic architecture of short stamen number, which has limited effects on seed set in the highly selfing plant *Arabidopsis thaliana*. We used QTL mapping to find loci affecting stamen number and other reproductive traits correlated with stamen number in nature. The same genomic region affected stamen number and other reproductive traits in one of our two replicate mapping populations, but not the other. The patterns of QTL co-localization are thus cross-specific.

## Introduction

Lineages across the tree of life have lost traits over macroevolutionary timescales, often when the function of the trait is lost or reduced. Widespread trait loss includes the loss of roots in aquatic ferns, the loss of teeth in birds, and the loss of tails in apes (Louchart and Viriot 2011; Ware et al. 2023; Xia et al. 2024). These textbook examples of trait loss reflect differences between species or higher taxa that occur over long evolutionary timescales. When intraspecific genetic variation exists for trait presence and absence, then processes like drift, selection for or against trait loss, and selection on genetically correlated traits will influence which populations lose the trait and which maintain it (Shapiro et al. 2004; Coyle et al. 2007; McGaugh et al. 2014; Cabin et al. 2022; Moran et al. 2023). By leveraging such intraspecific genetic variation for trait loss, we can investigate trait loss as a microevolutionary process which, over long timescales, may give rise to the morphological diversity we see among lineages.

Pleiotropy can cause correlated divergence of multiple traits. In wild radish (*Raphanus raphanistrum*), pleiotropy or tight linkage underlies a positive genetic correlation between two floral morphological traits, filament length and corolla tube length (Conner 2002), which reduced the response to artificial selection for longer filaments and shorter corolla tubes (Conner et al. 2011). Such pleiotropy may contribute to correlations between the sizes of different floral organs, although floral traits are generally only moderately correlated (Ordano et al. 2008; Conner et al. 2014). QTLs for multiple floral traits frequently overlap, suggesting a possible pleiotropic architecture (Juenger et al. 2000; Juenger et al. 2005; Slotte et al. 2012; Chen et al. 2025). The extent to which pleiotropic genes underlie trait correlations in nature is an open question in evolutionary biology.

Molecular genetics has uncovered numerous examples of genes that when perturbed in the lab, influence multiple floral traits. Rice (*Oryza sativa*) with mutations in *LONELY GUY1* has few to no stamens and pistils and fewer panicle branches (Kurakawa et al. 2007). Overexpression of *DREB2C* in *Arabidopsis thaliana* leads to delayed flowering and shorter stamen length (Song et al. 2022), and the E3 ubiquitin ligase *BIG BROTHER* pleiotropically affects the sizes of leaves and multiple floral organs by regulating cell division (Disch et al. 2006). These and numerous other molecular genetic experiments (reviewed in Auge et al. 2019) suggest that it is common for genes to function in multiple floral development pathways. However, whether natural variation exists in these pleiotropic genes, and whether such variation drives trait correlations in nature, remains relatively underexplored.

There are few cases where the molecular identity of genes pleiotropically underlying trait correlations in nature is known. In *Arabidopsis thaliana*, the gene *FRIGIDA* influences both water use efficiency and flowering time, contributing to a positive correlation between flowering time and water use efficiency (McKay et al. 2003; Lovell et al. 2013). In cavefish (*Astyanax mexicanus*), deletions in the gene *oca2* led to reduced pigmentation and reduced sleep compared to surface fish, suggesting that pigmentation loss may have evolved due to selection for reduced sleep (O’Gorman et al. 2021). Determining the genes responsible for quantitative trait variation is critical; without identifying genes, perturbing them through gene editing, and measuring multiple phenotypes, it is difficult to distinguish pleiotropy from linkage in genetic correlations (Saltz et al. 2017). Indeed, multiple morphological traits mapped to an overlapping genomic region in *Mimulus guttatus* and were highly correlated in recombinant mapping populations, suggesting that a pleiotropic gene may underlie multivariate trait divergence (Hall et al. 2006). However, later analysis revealed that the QTL corresponded to a large chromosomal inversion containing hundreds of physically linked genes (Lowry and Willis 2010). Determining the molecular identity of the genes underlying trait correlations (or individual traits, if not actually pleiotropic) offers substantial insight for the evolutionary mechanisms underlying phenotypic diversity.

The production of flowers bearing four long (medial) and two short (lateral) stamens is a diagnostic trait for the Brassicaceae family (Zomlefer 1994). The two stamen lengths improve pollination in outcrossing weedy radish (Waterman et al. 2025). However, variation in the production of the short stamens exists between Brassicaceae species (Endress 1992) and within the self-fertilizing species *Cardamine hirsuta* and *Arabidopsis thaliana* (Hay et al. 2014; Royer et al. 2016). In the latter species, the long stamen anthers are responsible for most or all self-pollination because the short stamen anthers are typically well below the stigma, and removing the short stamens does not reduce seed production significantly (Royer et al. 2016). Consistent with short stamens being non-essential for seed production, some *A. thaliana* individuals frequently produce flowers with 0 or 1 short stamens. Short stamen number is therefore an ideal trait to understand how populations lose traits with reduced function as a microevolutionary process, ultimately producing morphological diversity across the tree of life.

Multiple traits, including short stamen number, show latitudinal and altitudinal clines in *A. thaliana*. Short stamen loss is more common in *A. thaliana* at low elevations and low latitudes and is correlated with early flowering time and more ovules per flower (Royer et al. 2016; Buysse et al. 2025). If pleiotropic or tightly linked loci underlie variation in a trait under selection *and* stamen number, then geographic patterns of stamen number variation may reflect a correlated response to selection on the other trait, rather than the direct costs and benefits of short stamen number for plant fitness. In the Belm-12 x Roda-47 RILs constructed by crossing parents at the extremes of a latitudinal gradient, flowering time and short stamen number both map to a region of chromosome 5 that includes the flowering time regulator *FLC* (Dittmar et al. 2014; Royer et al. 2016), offering pleiotropy or linkage as a plausible mechanism for the correlation of these traits. Identifying the underlying gene or genes within this QTL is a key step for understanding the roles of linkage and pleiotropy in short stamen number evolution.

Determining whether QTLs affecting other traits repeatedly co-localize with stamen number QTLs is a key strategy for investigating the repeatability of trait loss genetic architecture. Specifically, if the same gene or QTL region affect both stamen loss and another trait, then such genetic architecture would offer a mechanism for linkage or pleiotropy to drive stamen number evolution. Given that different QTLs are commonly identified in different mapping populations of the same species (Dilda and Mackay 2002; Symonds et al. 2005; Kumar et al. 2007), patterns of overlapping QTLs that are shared among crosses will be more informative for understanding species-wide patterns of correlated evolution. Here, we used the Tsu-1 x Kas-1 RIL mapping population to identify QTLs associated with short stamen loss and ovule number in *A. thaliana*. We reanalyzed stamen number QTLs in the previously studied Belm-12 x Roda-47 RIL mapping population (Royer et al., 2016) and identified ovule number QTLs to determine whether QTLs for stamen number consistently co-localize between replicated mapping populations. Previous studies showed different sets of flowering time loci in these two sets of RILs (Lovell et al. 2013; Dittmar et al. 2014), and we repeated these analyses to verify that they reflect different genetic architectures rather than differences in QTL analysis pipelines. To select candidate genes affecting stamen number, we identified genomic variants within the overlapping chromosome 5 QTL region that are shared by the stamen loss parents of each cross. We phenotyped T-DNA insertion mutants in a Col-0 background for 13 candidate genes to test whether these knockouts have fewer short stamens than wild-type Col-0. Taken together, our investigation reveals that the genetic architecture of short stamen number is partially conserved between replicate mapping populations, and that co-localized QTLs affect short stamen number, flowering time, and ovule number in a genotype-specific manner.

## Materials and methods

### Plant Material and Growth Conditions

RILs were originally generated by crossing Tsu-1 (CS1640) and Kas-1 (CS903) reciprocally, obtaining the F2 progeny, and self-fertilizing to reach an F9 generation (McKay et al. 2008). We obtained these 341 RILs from the Arabidopsis Biological Resource Center (https://abrc.osu.edu/, ABRC). The Tsu-1 and Kas-1 parents were originally selected for the study of drought adaptation since they were sourced from habitats with contrasting water availability (Tsushima, Japan is wet throughout the growing season and Kashmir, India has little precipitation during the growing season; see Table 1, McKay et al. 2008). We selected these RILs for QTL mapping since the parents had divergent short stamen number phenotypes in preliminary observations and are distant longitudinally, rather than the latitudinal gradient spanned by the sites of origin for Belm-12 and Roda-47. The Belm-12 x Roda-47 RILs were originally generated for the study of local adaptation to Sweden and Italy (Ågren et al. 2013), with the Belm-12 parent sourced from Castelnuovo di Porto in central Italy and the Roda-47 parent sourced from Rödåsen, Sweden (Table 1). Belm-12 and Roda-47 were both sampled from populations that live as winter annuals on rocky slopes near forests, but they face contrasting water regimes, with 817 mm annual precipitation at the Italian site and 653 mm annual precipitation at the Swedish site (Ågren and Schemske 2012).

**Table 1.** Accessions used for QTL mapping in this study.

| Name | ABRC Accession | Source | Purpose in this study |
|---|---|---|---|
| Belm-12 | CS98761 | Parental line sampled from Castelnuovo di Porto, Italy, at 42.12°N, 12.48°E (Ågren and Schemske 2012). | Stamen loss, early flowering parent in Belm-12 x Roda-47 RILs |
| Roda-47 | CS98762 | Parental line sampled from Rödåsen, Sweden, at 62.80°N, 18.20°E (Ågren and Schemske 2012). | Stamen retaining, late flowering parent in Belm-12 x Roda-47 RILs |
| Belm-12 x Roda-47 RILs | CS98760 | Set of 544 RILs generated by crossing Belm-12 x Roda-47, selfing that F1 to produce F2’s, and propagating those F2’s through 9 generations of single-seed descent (Ågren et al. 2013). | QTL mapping |
| Tsu-1 | CS1640 | Parental line sampled from Tsushima, Japan, at 34.5°N, 136.3°E (McKay et al. 2003). | Stamen retaining, early flowering parent in Tsu-1 x Kas-1 RILs |
| Kas-1 | CS903 | Parental line sampled from Kashmir, India, at 35°N, 77°E (McKay et al. 2003). | Stamen loss, late flowering parent in Tsu-1 x Kas-1 RILs |
| Tsu-1 x Kas-1 RILs | CS96685-CS97025 | Set of 346 RILs generated by crossing Tsu-1 x Kas-1, selfing that F1 to produce F2’s, and propagating those F2’s through 9 generations of single-seed descent (McKay et al. 2008) | QTL mapping |

To characterize parental trait divergence and perform QTL mapping, ten individuals from each of the Tsu-1 and Kas-1 parental accessions and two individuals from each Tsu-1 x Kas-1 RIL were grown and phenotyped. The first temporal block of one plant from each RIL and five plants from each parental accession was sown in moist Metro-Mix, stratified at 4°C for 6 days, and transferred to a greenhouse at Kellogg Biological Station (Hickory Corners, Michigan, USA) in July 2014 that received approximately 15 hours of light per day, with 22°C day and 19°C night temperatures. A second temporal block with the same composition was grown under the same conditions in August 2014, except with a day temperature of 24°C.

Belm-12 x Roda-47 RIL growth conditions are described in Royer et al. (2016) for stamen and ovule number, and in Dittmar et al. (2014) for flowering time in the simulated Italy chamber. We note that Dittmar et al. (2014) used growth chamber conditions mimicking environmental conditions in Italy, which included a simulated winter.

T-DNA insertion mutant lines in the Col-0 genetic background were ordered from ABRC (accessions and genotyping primers listed in Supplementary Table 1 in File S1). Wild-type (Col-0) and mutant plants were grown in a growth chamber with a 22°C day and night temperature and light intensity of 120-150 µmol m^-2^ s^-1^ for 16 hours each day, with some lines phenotyped in 2022 and additional lines in 2025 (Supplementary Table 2 in File S1). Individual plants were grown in 2.5-inch square pots and bottom watered three times weekly. Trays of plants were rotated and moved within the growth chamber 1-2 times per week, and mutant lines were compared to Col-0 individuals randomized within their tray.

### Flowering Time, Stamen Number, and Ovule Number RIL Phenotyping

We recorded the germination date and day of first flower to calculate flowering time from germination. We collected freshly opened flowers and stored them in 70% alcohol on each of three separate dates occurring across the flowering period, recording the flower positions collected and the collection date for each timepoint. We recorded the number of long and short stamens for each flower by viewing with a dissecting scope from a mean of 5.8 flowers per plant. Ovule counts followed Royer et al. (2016). Briefly, we placed the pistil of one flower from each of the three collection dates for a total of three flowers per plant on a glass slide, applied a drop of blue food coloring and a coverslip, and counted ovules with a hand tally counter. Ovule counts for Belm-12 x Roda-47 RILs were performed using flowers collected in Royer et al. (2016). We estimated mean trait values for each RIL as the least squares means (LSM) from a linear model in JMP (JMP, Version 15. SAS Institute Inc., Cary, NC, 1989-2021) that included flower position within an inflorescence, type of inflorescence (main or side inflorescence), block, and collection date. LSMs of each trait were highly correlated with raw means for all traits (short stamen number correlation coefficient = 0.994; flowering time correlation coefficient = 0.999; ovule number correlation coefficient = 0.864), so raw trait means across the two plants measured for each RIL were used in downstream QTL mapping to maintain a consistent analytical pipeline with Belm-12 x Roda-47 RILs.

Stamen number data for the Belm-12 x Roda-47 RILs were obtained from a previous study (Royer et al. 2016). Flowering time data were obtained a previous study (Dittmar et al. 2014) for for Belm-12 x Roda-47 RILs. Since many RILs did not flower in the chamber simulating the Sweden environment in Dittmar et al. (2014), we only used flowering time data from the chamber simulating the Italy environment.

### Analysis of RIL Correlations

We estimated genetic correlations in the RILs as Pearson correlation coefficients between RIL mean phenotypes using the R package *corrtable* (version 0.1.1). Trait correlations in RIL or F2 mapping populations are evidence of linkage or pleiotropy; however, mapping populations display variation in the percentage of the genome that is contributed by each parent (Frisch and Melchinger 2007). If RIL parents differ in two traits, then some RILs may resemble one parent or the other for both traits due to solely to inheriting a high proportion of that parent’s genome, including loci that are neither pleiotropic nor in linkage disequilibrium in the parental populations. Thus, the covariances between traits in RILs would include many loci that only affect one trait that are in linkage equilibrium in the parental populations, but the admixture between the divergent parents and variation in parental genome content creates novel linkage disequilibrium between these independent loci.

To understand the role of parental genome contribution, linkage, and pleiotropy in observed trait correlations, we designed an approach to approximate the fraction of Tsu-1 genome in the Tsu-1 x Kas-1 RILs as the number of homozygous Tsu-1 markers divided by the total number of markers (excluding missing genotypes), and the fraction of Roda-47 genome in the Belm-12 x Roda-47 RILs as the number of homozygous Roda-47 markers divided by the total number of markers (excluding missing genotypes). We then used the R package *ppcor* (version 1.1) to test for partial correlations between traits, controlling for the fraction of Tsu-1 genome in Tsu-1 x Kas-1 RILs and the fraction of Roda-47 genome in the Belm-12 x Roda-47 RILs (but not removing effects of other phenotypic traits). We considered the alternative approach of using the relatedness matrix to estimate the genetic correlations in the RILs, but this estimate would include the novel linkage disequilibrium created by admixture and thus would not estimate correlations caused only by pleiotropy or tight linkage.

### QTL Mapping

Linkage maps and genotypes for the RILs were generated previously. Briefly, the Tsu-1 x Kas-1 RILs were genotyped at 450 markers, consisting of single nucleotide polymorphisms (SNPs) and simple sequence length polymorphisms (Lovell et al. 2015). The Belm-12 x Roda-47 RILs were genotyped at 348 SNP markers (Ågren et al. 2013).

We used the R/qtl package version 1.70 (Broman and Sen 2009) to carry out QTL analyses in R (version 4.1.1). To detect large-effect QTLs, we first performed one-dimensional QTL mapping with Haley-Knot regression and 10,000 permutations to set a LOD threshold with significance level set at 0.05. To determine significance levels for downstream multi-QTL mapping, we then performed two-dimensional analyses allowing for pairwise interactions, again with Haley-Knot regression and 10,000 permutations, and used LOD thresholds at significance levels of 0.05 and 0.01 in multi-QTL modeling. To improve detection of small-effect QTLs by controlling for other detected QTLs, we constructed multi-QTL models at each significance level using automated computation with the *stepwise* function in R/qtl, allowing for epistasis and examining models produced with *max.qtl* parameters between 5-15. The *stepwise* function compares all models with between 1 and *max.qtl* QTL, so the *max.qtl* parameter constrains the search space of *stepwise.* The significance threshold of 0.01 identified the same significant QTL for *max.qtl* values between 6-15, whereas the significance threshold of 0.05 found more QTLs as *max.qtl* increased. Since the significance threshold of 0.01 led to a position and number of QTLs that was robust to changes in *max.qtl,* the multi-QTL model reported here used a significance level of 0.01 and *max.qtl* of 15. To verify that QTLs found in our analysis are robust to the non-normal distribution of stamen number data, we performed the QTL analysis workflow described above for quantile-normalized short stamen number calculated using the bestNormalize R package (version 1.9.1).

Stepwise modeling of raw short stamen number in Tsu-1 x Kas-1 RILs initially identified a pair of linked QTLs on chromosome 5 at 2.8 and 9.8 cM. Linked QTLs are challenging to distinguish from a single, large QTL due to limited recombination between them and the importance of precisely locating the ends of QTL regions. Since only one QTL was identified on chromosome 5 with quantile-normalized data, and map position in the Tsu-1 x Kas-1 linkage map does not always increase with increasing base pair position in the Col-0 reference genome, we removed the smaller effect QTL at 9.8 cM using the *dropfromqtl* function and refined the positions of remaining QTLs using *refineqtl*.

To determine the genetic architecture of ovule number, we performed QTL analysis as described above on ovule number per flower for both crosses. We mapped the genetic architecture of flowering time in Tsu-1 x Kas-1 RILs using flowering time data collected in this study and the same workflow. For Belm-12 x Roda-47 RILs, we used data from Dittmar et al. (2014) collected in the chamber simulating the Italy environment (since the Sweden environment had a high rate of missing data) and followed the same workflow described for the other traits, to ensure that any differences between QTLs detected in the two mapping populations are due to biological differences rather than different analytical choices between previous flowering time publications (Lovell et al. 2013; Dittmar et al. 2014).

To enable comparison between stamen number QTL mapping using Tsu-1 x Kas-1 RILs and the previously characterized Belm-12 x Roda-47 RILs (Royer et al., 2016), we repeated QTL mapping of stamen loss for the Belm-12 x Roda-47 RILs using the same analysis parameters that were used above for Tsu-1 x Kas-1 RILs.

### Analysis of Putative Linked QTLs

Multi-QTL analysis identified two tightly linked QTLs on each of chromosomes 1 and 5 affecting short stamen number in Tsu-1 x Kas-1 RILs, even at a somewhat stringent significance level of 0.01. To further test whether there are two linked QTLs on these chromosomes, we excluded RILs with imputed genotypes at either of the markers at the peaks of chromosome 1 QTLs (1@80, 1@108) from the dataset and fit ANOVAs testing for the effect of genotype class (Kas-1 at both loci, Kas-1 at 1@80 and Tsu-1 at 1@108, Tsu-1 at 1@80 and Kas-1 and 1@108, and Tsu-1 at both loci) on RIL mean short stamen number. We tested for significant short stamen number differences between genotype classes using Tukey’s Honestly Significant Difference test. We performed the same analysis workflow for the two putative linked QTLs on chromosome 5, 5@2.8 and 5@9.8.

### Tests for Linked or Pleiotropic QTLs on Chromosome 5

QTLs affecting short stamen number, ovule number, and flowering time all mapped between 7.7-10 cM on chromosome 5 in the Belm-12 x Roda-47 cross. To test whether separate linked QTLs likely underlie these traits (as opposed to one pleiotropic QTL), we used the R package *qtl2pleio* (version 1.4.4) and R version 4.6.1 to test the null hypothesis that a single pleiotropic QTL underlies the traits of interest, for each pairwise combination of the three traits (Boehm et al. 2019). We calculated the leave-one-chromosome-out kinship matrix with R/qtl2 (version 0.46, Broman et al. 2019). We used the scan_pvl() and calc_lrt_tib() functions of *qtl2pleio* to estimate the likelihood ratio test statistic from two-dimensional scans. We performed 1000 parametric bootstrap replicates using the boot_pvl() function, covering the 1.5 LOD intervals of the two QTLs being examined as well as the marker where the likelihood of pleiotropy is highest. Reported *P-*values correspond to the fraction of bootstrap replicates for which the log10 likelihood ratio test statistic was greater than the likelihood ratio test statistic from calc_lrt_tib(). *P-*values less than 0.05 let us reject the null hypothesis that a single pleiotropic locus underlies the two traits examined.

### Genomic Analysis of RIL Parents

To identify candidate genes for stamen loss, we analyzed parental sequences within the QTL region on chromosome 5 identified in both the Tsu-1 x Kas-1 (this study) and Belm-12 x Roda-47 RILs (Royer et al., 2016). We extracted genomic DNA from leaf tissue of Tsu-1 (CS1640), Kas-1 (CS903), Belm-12 (CS98761), and Roda-47 (CS98762) using a modified CTAB method. Whole genome sequencing libraries were prepared at the MSU Genomics Core using the Roche KAPA HyperPrep DNA Library Preparation Kit with KAPA Unique Dual Index (UDI) adapters. Sequencing was performed in a ×2×150bp paired end format using a NovaSeq 6000 v1.5 300 cycle reagent kit on an S4 flow cell. Approximately 230 million paired-end reads were generated for each accession. We trimmed reads using fastp version 1.0.1 with default settings (Chen et al. 2018), aligned reads to the TAIR10 reference genome (Lamesch et al. 2012) using bwa-mem2 (Vasimuddin et al. 2019), and called variants using GATK version 4.6.2.0 (McKenna et al. 2010). We used VCFtools (version 0.1.16, Danecek et al. 2011) to filter for variants with a quality score of 30 or greater and coverage between 10 and 250x.

To select candidate genes that may underlie this shared chromosome 5 QTL, we used BCFtools (version 1.22) to identify homozygous variants that are shared by the stamen-loss parents Kas-1 and Belm-12 where the stamen-retaining parents Tsu-1 and Roda-47 have the reference allele (Danecek et al. 2021). We predicted the effects of such variants on gene function using snpEff version 4.3p (Cingolani et al. 2012) and prioritized candidate genes in and near the shared QTL region (roughly 1.5-3 Mb on chromosome 5) that had mutations shared by stamen loss parents that were expected to affect gene function. We performed functional analysis on such genes that were expressed in floral tissues (Klepikova et al. 2016) and have available T-DNA insertion mutants.

### T-DNA Insertion Mutant Genotyping and Phenotyping

We tested a total of 23 mutant lines of 13 genes (Table 7). We ordered T-DNA insertion lines from ABRC, extracted genomic DNA (Hu and Lagarias 2020), and genotyped mutant lines using primers binding and flanking the insertion site (Supplementary Table 1 in File S1, primers designed with SALK T-DNA Primer Design Tool, http://signal.salk.edu/tdnaprimers.2.html) to identify homozygous mutant lines for phenotyping. When phenotyping each line, 4-5 homozygous mutant plants and 4-5 wild-type (Col-0) plants were randomized in the same tray, with 1-2 mutant lines compared to wild-type plants within the same tray. We recorded short stamen number on flowers collected over developmental time and viewed with a dissecting scope, aiming to collect the 7^th^-9^th^, 17^th^-19^th^, and 27^th^-29^th^ flowers at anthesis from the main inflorescence on each plant so that phenotyping spanned plant development. We compared short stamen number of homozygous T-DNA insertional mutants to Col-0 plants grown in the same tray using ANOVAs in R (version 4.1.1).

## Results

### Phenotypic Variation in RILs

The parents of both sets of RILs differ significantly in short stamen number. For Tsu-1 x Kas-1 RILs, Kas-1 has short stamen loss (mean of 0.68 short stamens per flower) whereas Tsu-1 has a mean of 1.99 short stamens per flower (*P* < 0.0001, Figure 1a, Supplementary Figure 1a in File S2). As shown previously (Royer et al. 2016), Belm-12 produces 0.92 short stamens per flower on average and Roda-47 produces 1.92 (*P* < 0.0001, Figure 1b, Supplementary Figure 1b in File S2). For both mapping populations, most RILs have phenotypes between the parental means, as the maximum value of the trait is 2 short stamens. We observe some transgressive segregation for loss where RILs have fewer short stamens than the stamen-loss parent (Figure 1a, 1b, Supplementary Figure 1 in File S2). The Tsu-1 and Kas-1 accessions have similar ovule number (effect = −0.60, *P* = 0.79), while Belm-12 has more ovules per flower than Roda-47 (effect = - 5.95 ovules, *P* = 0.0005). Both sets of RILs showed transgressive segregation well outside the range of parental phenotypes (Figure 1c, 1d). The parental accessions show contrasting combinations of flowering time and stamen number phenotypes. Tsu-1 flowers earlier than Kas-1 yet retains short stamens (Figure 1a, 1e). In contrast, the stamen loss parent Belm-12 flowers earlier than stamen-retaining parent Roda-47 (Figure 1b, 1f), consistent with their sampling at extremes of a latitudinal cline.

**Figure 1.**
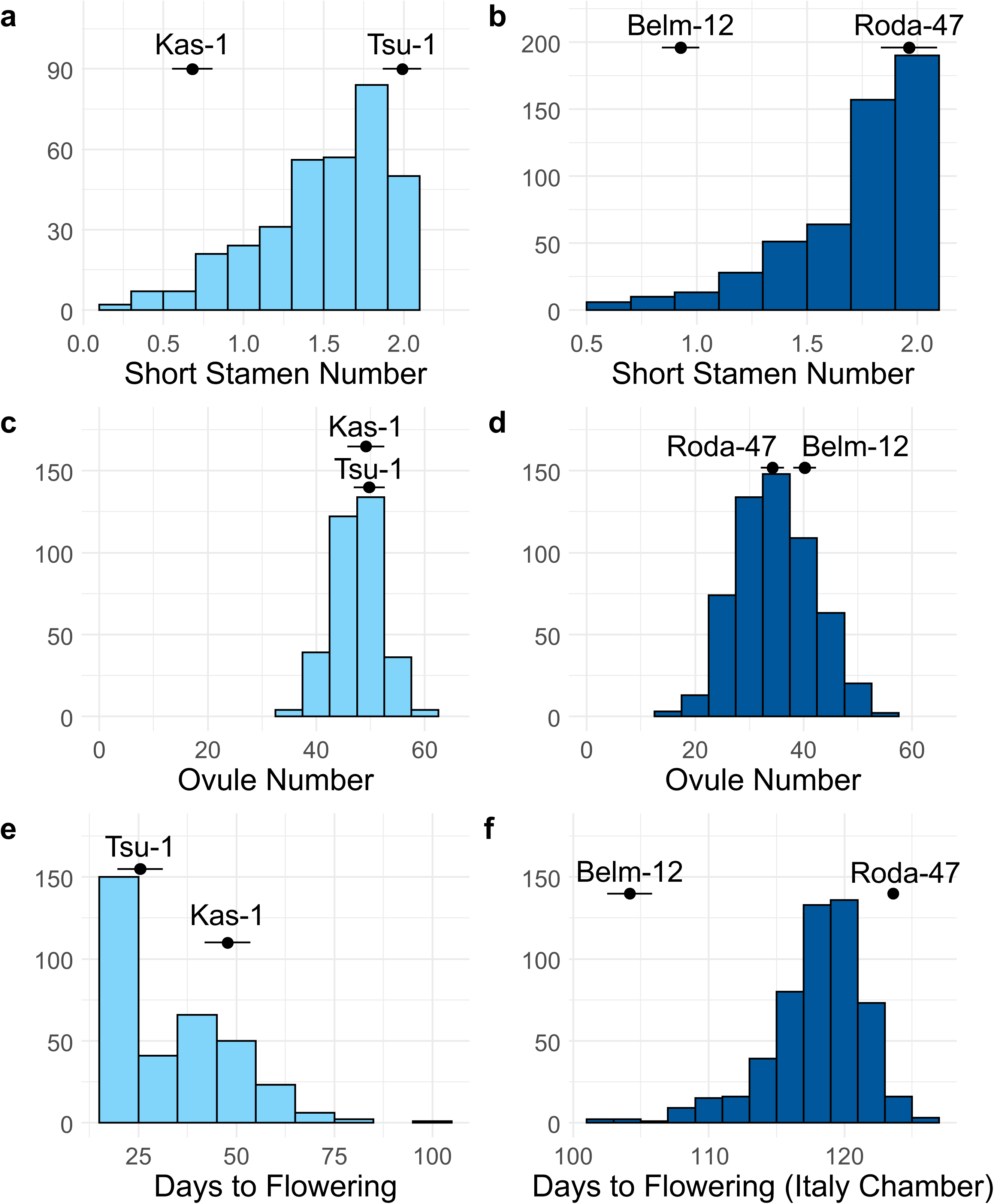
Trait distributions in RIL mapping populations, with parental accession mean phenotypes overlaid. Error bars indicate 95% confidence intervals. (a) Mean short stamen number for Tsu-1 x Kas-1 RILs, from this study. (b) Mean short stamen number for Belm-12 x Roda-47 RILs, from Royer et al., 2016. (c) Mean ovule number for Tsu-1 x Kas-1 RILs, from this study. (d) Mean ovule number for Belm-12 x Roda-47 RILs, from this study. (e) Days to flowering for Tsu-1 x Kas-1 RILs, from this study. (f) Days to flowering for Belm-12 x Roda-47 RILs, from Dittmar et al. (2014). Note different x-axes for (e) and (f). Alt text: Histograms showing phenotypic distributions in both sets of RILs, with trait means at 95% confidence intervals for the parental accessions. Tsu-1 x Kas-1 RILs are shown on the left and Belm-12 x Roda-47 RILs on the right. (a) Distribution of mean short stamen number in Tsu-1 x Kas-1 RILs. RIL phenotypes are between 0 and 2 short stamens, with most of the distribution close to 2. Tsu-1 has a mean short stamen number of 2 and Kas-1 has a mean short stamen number of 0.68. (b) Distribution of mean short stamen number in Belm-12 x Roda-47 RILs. RIL phenotypes are between 0 and 2 short stamens, with most of the distribution close to 2. Roda-47 has a mean short stamen number of 1.92 and Kas-1 has a mean short stamen number of 0.92. (c) Distribution of mean ovule number in Tsu-1 x Kas-1 RILs. RIL phenotypes are between 30-60 ovules per flower, with parental phenotypes at the center of the distribution. (d) Distribution of mean ovule number in Belm-12 x Roda-47 RILs. RIL phenotypes are between 10-60 ovules, with Roda-47 producing about 35 and Belm-12 producing about 40 ovules on average. (e) Distribution of flowering time in Tsu-1 x Kas-1 RILs. RIL phenotypes are 100 or fewer days to flowering with the most common flowering time under 25 days. Tsu-1 took about 25 days to flower and Kas-1 took about 48 days. (f) Distribution of flowering time in Belm-12 x Roda-47 RILs in the Italy chamber environment from Dittmar et al. (2014). RIL phenotypes are between 100-140 days, with Belm-12 flowering in 104 days and Roda-47 flowering in 124 days.

### Trait Correlations with Short Stamen Number

Neither flowering time nor ovule count traits were correlated with short stamen number in the Tsu-1 x Kas-1 RILs, indicating that loci underlying stamen number variation are not strongly linked nor pleiotropic in this cross. In contrast, in the Belm-12 x Roda-47 RILs, flowering time was positively correlated with short stamen number, and ovule count was negatively correlated with short stamen number (Table 2). Thus, co-localization of flowering time or ovule number QTLs may drive these trait correlations with stamen number in Belm-12 x Roda-47 RILs. However, the lack of these correlations in Tsu-1 x Kas-1 RILs indicates that these correlations are specific to certain genetic architectures of flowering time and ovule number.

**Table 2.**
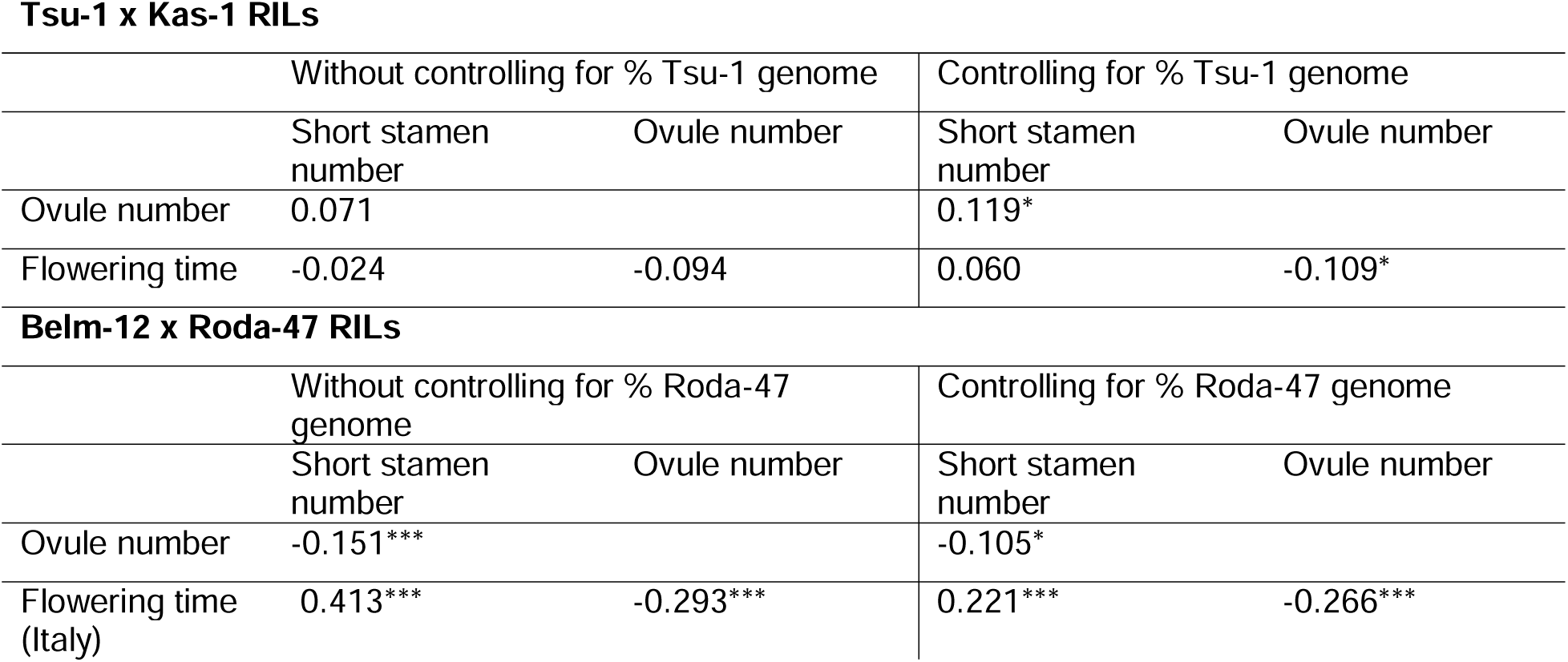
Pairwise correlations between short stamen number, days to flowering, and mean ovule count in Tsu-1 x Kas-1 RILs (this study), and stamen number (Royer et al. 2016), ovule number (this study), and flowering time (Dittmar et al. 2014) in Belm-12 x Roda-47 RILs. Significance levels are indicated as * (p < 0.05), ** (p < 0.01), and *** (p < 0.001).

| **Tsu-1 x Kas-1 RILs** |  |  |  |  |
|---|---|---|---|---|
|  | Without controlling for % Tsu-1 genome |  | Controlling for % Tsu-1 genome |  |
|  | Short stamen number | Ovule number | Short stamen number | Ovule number |
| Ovule number | 0.071 |  | 0.119* |  |
| Flowering time | -0.024 | -0.094 | 0.060 | -0.109* |
| **Belm-12 x Roda-47 RILs** |  |  |  |  |
|  | Without controlling for % Roda-47 genome |  | Controlling for % Roda-47 genome |  |
|  | Short stamen number | Ovule number | Short stamen number | Ovule number |
| Ovule number | -0.151*** |  | -0.105* |  |
| Flowering time (Italy) | 0.413*** | -0.293*** | 0.221*** | -0.266*** |

### Parental Genome Contributions to RILs

Both sets of RILs displayed large variation in the proportion of the genome contributed by each of the parents. Estimated as the proportion of markers corresponding to each parental genotype, the Tsu-1 x Kas-1 RILs had 12.7 – 81.0% of the genome contributed by the Tsu-1 parent, while the Belm-12 x Roda-47 RILs had 0 – 100% of the genome contributed by the Roda-47 parent (Supplementary Figure 2 in File S2). The extremes of the distribution for the Roda-47 parental genome contribution seem unlikely, as all RILs are descended from a Belm-12 x Roda-47 F1 individual; however, only 8 out of the 544 RILs had a Roda-47 genomic contribution of 0 or 100%. We speculate that these RILs may have very small regions of Belm-12 introgression between markers that is not detectable with our approximation, or contamination with parental seed. Excluding these eight individuals, the Roda-47 parental genome contribution ranges from 13.0 – 92.0%.

Controlling for the fraction of RIL genome contributed by each parent led to changes in pairwise correlation coefficients. For Belm-12 x Roda-47 RILs, traits that were correlated without controlling for the fraction of RIL genome from Roda-47 remained significant in the partial correlation analysis, but were weaker in magnitude. For example, flowering time in the Italy chamber was correlated with stamen number (r = 0.412, p < 0.001), and was more weakly correlated after controlling for the Roda-47 genomic contribution (r = 0.221, p < 0.001; Table 2). The uncontrolled pairwise correlation between flowering time and stamen number in Belm-12 x Roda-47 RILs may reflect linked or pleiotropic loci underlying the two traits as well as patterns of unlinked divergence in the two traits between parents (that is, RILs that inherit a high proportion of the Roda-47 genome are likely to resemble Roda-47 for multiple traits, which may contribute to pairwise trait correlation in RILs). Correlations were already low in the Tsu-1 x Kas-1 RILs, perhaps because these parents were not divergent for ovule number and were less divergent for flowering time than the Belm-12 x Roda-47 RILs, so little linkage disequilibrium for these traits was created by crossing the Tsu-1 and Kas-1 parents.

### Genetic Architecture of Short Stamen Number

In the Tsu-1 x Kas-1 RILs, we identified five QTLs with significant additive effects and one significant pairwise interaction for short stamen loss, which together explained 61.4% of the variation in RIL mean short stamen number. These QTLs are located on chromosomes 1, 3, 4, and 5, with two linked QTLs detected on chromosome 1 (Table 3, Figure 2a).

**Figure 2.**
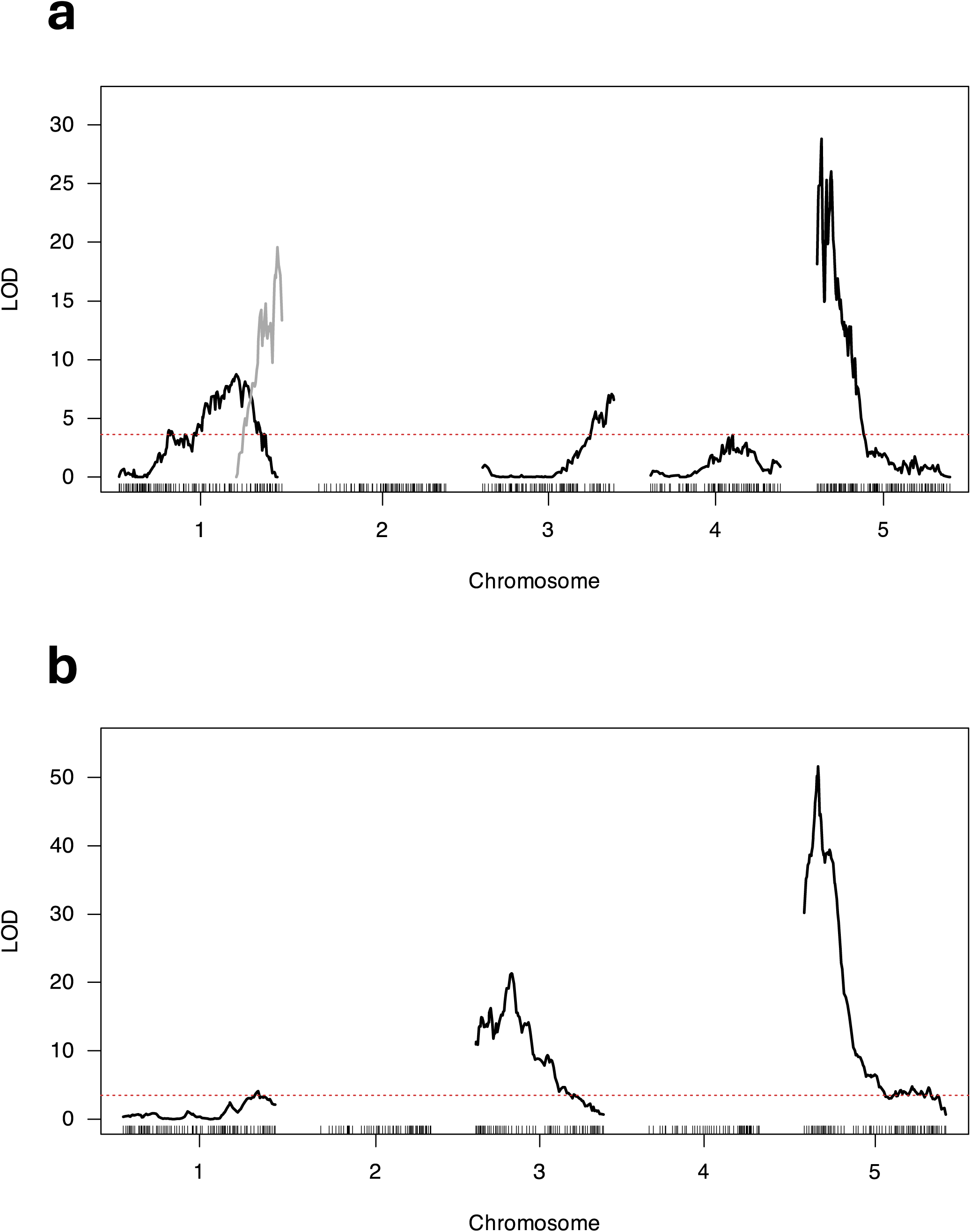
LOD plots showing multi-QTL mapping of short stamen number for Tsu-1 x Kas-1 (a) and Belm-12 x Roda-47 (b) mapping populations. Red dashed lines indicate thresholds from scantwo permutations at significance level of 0.01. Each trace indicates the LOD profile for a separate QTL, with gray and black used to help distinguish QTL traces on the same chromosome. Alt text: Graph with LOD scores on the y-axis and genetic map position for 5 chromosomes on the x-axis. (a) LOD traces for 5 short stamen number QTLs in Tsu-1 x Kas-1. The LOD peaks of QTLs on chromosomes 1, 3, and 5 are well above the LOD threshold (about 4) while the QTL on chromosome 4 barely reaches it.

**Figure 3.**
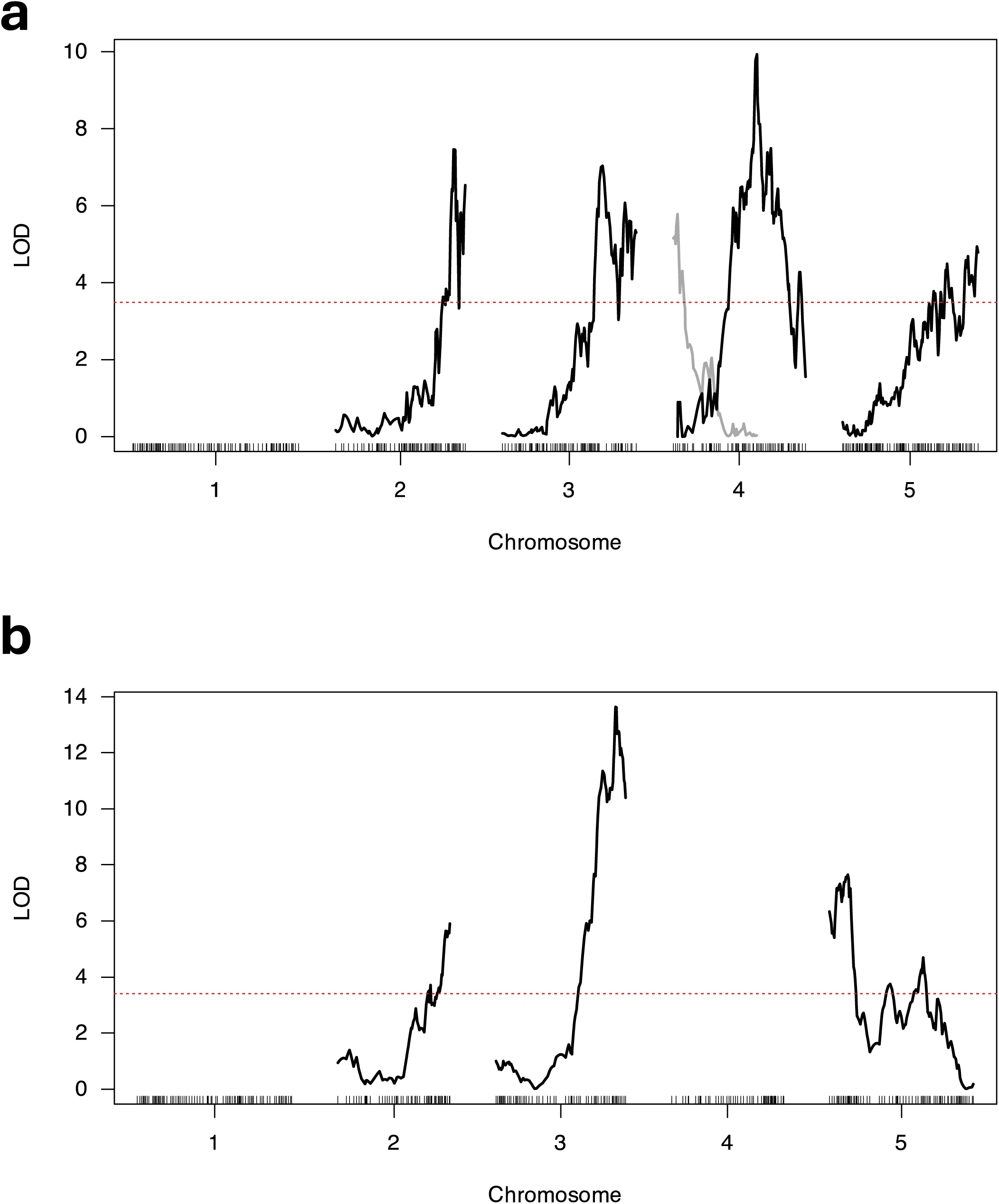
LOD plots showing multi-QTL mapping of ovule number for Tsu-1 x Kas-1 (a) and Belm-12 x Roda-47 (b) mapping populations. Red dashed lines indicate thresholds from scantwo permutations at significance level of 0.01. Each trace indicates the LOD profile for a separate QTL, with gray and black used to help distinguish QTL traces on the same chromosome (see Table 5). Alt text: Graph with LOD scores on the y-axis and genetic map position for 5 chromosomes on the x-axis. (a) LOD traces for 5 ovule number QTLs in Tsu-1 x Kas-1 on chromosomes 2, 3, 4, 4, and 5. (b) LOD traces for 5 ovule number QTLs in Belm-12 x Roda-47 on chromosomes 2, 3, and 5.

**Table 3.** Short stamen number QTL from both mapping populations. The additive effect on short stamen number for one allele from the stamen-retaining parent is given in a, %var is the percent of RIL variation explained by that QTL, and %div is percent of parental divergence explained, as 100 * 2a / (stamen retaining parent mean short stamen number – stamen loss parent mean short stamen number).

| Tsu-1 x Kas-1 RILs |  |  |  |  |  |  |  |  |
| QTL | Chr | Position (cM) | 1.5 LOD Interval (cM) | 1.5 LOD Interval (Mb) | LOD | a | %var | %div |
|---|---|---|---|---|---|---|---|---|
| 1@80.0 | 1 | 80 | 66.40 – 89 | 20.53 – 25.59 | 9.25 | 0.11 | 5.17 | 16.55 |
| 1@108.0 | 1 | 108 | 107.43 – 108.94 | 27.65 – 28.67 | 26.05 | 0.17 | 16.38 | 25.50 |
| 3@88.0 | 3 | 88 | 84 – 89.50 | 19.93 – 21.63 | 8.08 | -0.09 | 4.48 | -14.31 |
| 4@56.1 | 4 | 56.1 | 48.07 – 71 | 8.30 – 19.16 | 4.19 | 0.07 | 2.26 | 10.02 |
| 5@3.0 | 5 | 3.0 | 2.01 – 3.80 | 0.87 – 0.48 | 35.26 | 0.20 | 23.71 | 20.90 |
| Interaction 1@108.0:5@3.0 |  |  |  |  | 6.46 | -0.09 | 3.54 | -13.22 |
| Belm-12 x Roda-47 RILs |  |  |  |  |  |  |  |  |
| QTL | Chr | Position (cM) | 1.5 LOD Interval (cM) | 1.5 LOD Interval (Mb) | LOD | a | %var | %div |
| 1@74.4 | 1 | 74.44 | 67 - 82 | 25.03 – 29.38 | 4.05 | 0.05 | 2.1 | 8.78 |
| 3@19.7 | 3 | 19.71 | 18.03 - 22 | 7.91 – 8.50 | 21.3 | 0.09 | 11.96 | 18.12 |
| 5@7.7 | 5 | 7.65 | 6.68 - 8 | 2.41 – 2.99 | 51.63 | 0.18 | 33.39 | 34.44 |
| Interaction 3@19.7:5@7.7 |  |  |  |  | 9.32 | -0.07 | 4.95 | -13.77 |

In Belm-12 x Roda-47 RILs, we also recovered QTLs on chromosomes 1, 3, and 5, recapitulating the findings of Royer et al. (2016, Table 3, Figure 2b). The three QTLs and one pairwise interaction explain 42.54% of variation among Belm-12 x Roda-47 RILs. Both RIL sets had a large-effect QTL on chromosome 5, within 2 Mb of each other when compared using marker positions in the Col-0 reference genome (since the two crosses used different linkage maps). On chromosome 1, the 1.5-LOD support intervals in Belm-12 x Roda-47 RILs overlap with QTLs from the Tsu-1 x Kas-1 on chromosome 1 (Table 3, Figure 2). Notably, the chromosome 1 QTL in Belm-12 x Roda-47 RILs has a much smaller effect on short stamen number than either of the chromosome 1 QTLs in Tsu-1 x Kas-1 RILs (Table 3). The QTLs in Tsu-1 x Kas-1 chromosomes 3 and 4 are unique to the Tsu-1 x Kas-1 RILs.

For the Tsu-1 x Kas-1 RILs, QTL mapping results showed similarities and differences when using raw short stamen number or quantile-normalized short stamen number. In Tsu-1 x Kas-1, all of the QTLs recovered using raw short stamen number data were also recovered in the analysis with quantile-normalized short stamen number (Supplementary Table 3, Supplementary Figure 3 in File S2). We observed shifts in the position of QTL peaks and found a third QTL on chromosome 1 when using quantile-normalized short stamen number with Tsu-1 x Kas-1 RILs, but the 1.5 LOD intervals for these QTLs overlap between the two analyses. The pairwise interaction between QTLs on chromosomes 1 and 5 was consistent with and without quantile-normalized short stamen number. Finally, our initial analysis of raw short stamen number found two linked QTLs at 2.8 cM and 9.8 cM on chromosome 5 (not shown). We suspected that the large changes in LOD across chromosome 5 may be an artefact of possible errors in marker order (Supplementary Figure 5 in File S2) or the non-normal distribution of short stamen number. For this reason, we removed the QTL at 9.8 cM and Tsu-1 x Kas-1 RILs to have a single large-effect QTL on chromosome 5 at 3.0 cM.

For Belm-12 x Roda-47 RILs, we found identical positions of QTL peaks with and without quantile-normalization, with slight differences in 1.5 LOD intervals and effect sizes (Table 3, Supplementary Table 3 in File S2). These QTL positions are similar to those found by Royer et al. (2016). We recovered the pairwise interaction between QTLs on chromosomes 3 and 5, but not the marginally significant interaction reported in Royer et al. (2016) between QTLs on chromosomes 1 and 5. We note that we tested for interactions by allowing epistasis in our multi-QTL model, whereas Royer et al. (2016) tested for all possible pairwise interactions using ANOVAs after fitting the multi-QTL model.

### Genetic Architecture of Ovule Number

We identified 5 ovule number QTLs and 1 pairwise interaction in the Tsu-1 x Kas-1 RILs and 3 ovule number QTLs in the Belm-12 x Roda-47 RILs (Table 4). The QTLs on chromosomes 2 and 3 are shared between mapping populations. Although both sets of RILs had an ovule number QTL on chromosome 5, only the Belm-12 x Roda-47 QTL overlaps with the shared stamen number QTL region on chromosome 5 (Table 3, Table 4). The overlapping QTL regions for short stamen and ovule number may drive the correlation between these traits, which is observed in the Belm-12 x Roda-47 but not Tsu-1 x Kas-1 RILs (Table 2).

**Table 4.**
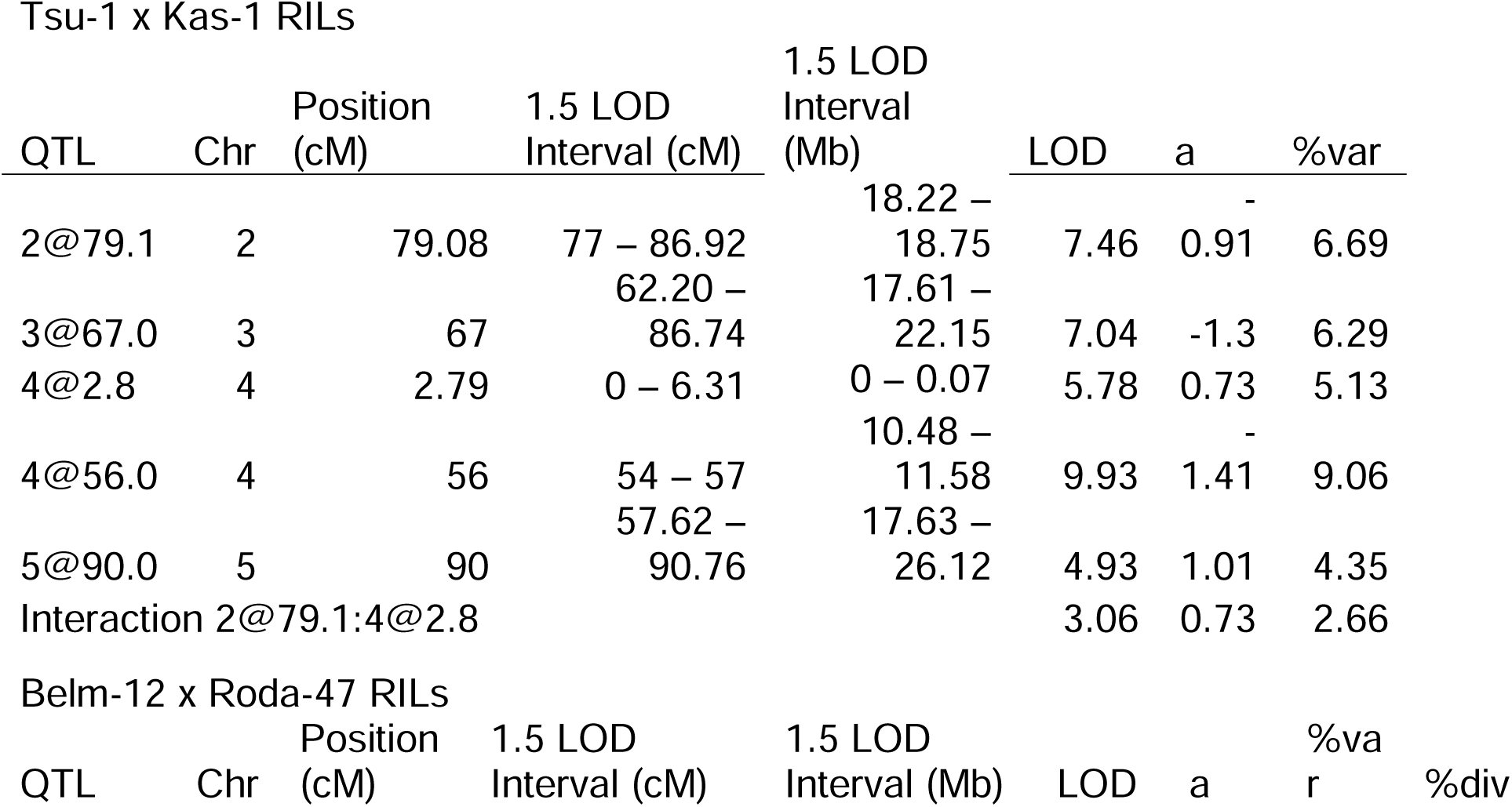
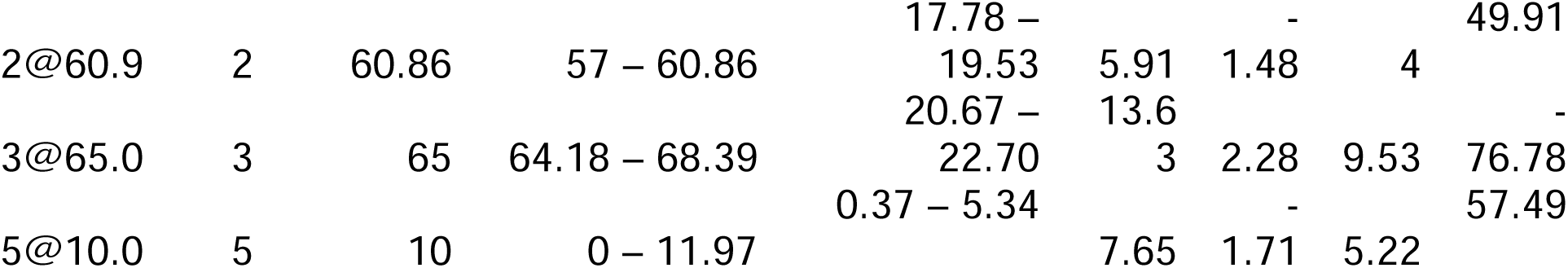
Ovule number QTL from both mapping populations. The additive effect on ovule number for one allele from the stamen-retaining parent is given in a, %var is the percent of RIL variation explained by that QTL, and %div is percent of parental divergence explained, as 100 * 2a / (stamen retaining parent mean ovule number – stamen loss parent mean ovule number).

| Tsu-1 x Kas-1 RILs |  |  |  |  |  |  |  |  |
|---|---|---|---|---|---|---|---|---|
| QTL | Chr | Position (cM) | 1.5 LOD Interval (cM) | 1.5 LOD Interval (Mb) | LOD | a | %var |  |
| 2@79.1 | 2 | 79.08 | 77 – 86.92 | 18.22 – 18.75 | 7.46 | -0.91 | 6.69 |  |
| 3@67.0 | 3 | 67 | 62.20 – 86.74 | 17.61 – 22.15 | 7.04 | -1.3 | 6.29 |  |
| 4@2.8 | 4 | 2.79 | 0 – 6.31 | 0 – 0.07 | 5.78 | 0.73 | 5.13 |  |
| 4@56.0 | 4 | 56 | 54 – 57 | 10.48 – 11.58 | 9.93 | -1.41 | 9.06 |  |
| 5@90.0 | 5 | 90 | 57.62 – 90.76 | 17.63 – 26.12 | 4.93 | 1.01 | 4.35 |  |
| Interaction 2@79.1:4@2.8 |  |  |  |  | 3.06 | 0.73 | 2.66 |  |
| Belm-12 x Roda-47 RILs |  |  |  |  |  |  |  |  |
| QTL | Chr | Position (cM) | 1.5 LOD Interval (cM) | 1.5 LOD Interval (Mb) | LOD | a | %var | %div |
| 2@60.9 | 2 | 60.86 | 57 – 60.86 | 17.78 – 19.53 | 5.91 | -1.48 | 4 | 49.91 |
| 3@65.0 | 3 | 65 | 64.18 – 68.39 | 20.67 – 22.70 | 13.63 | 2.28 | 9.53 | -76.78 |
| 5@10.0 | 5 | 10 | 0 – 11.97 | 0.37 – 5.34 | 7.65 | -1.71 | 5.22 | 57.49 |

### Genetic Architecture of Flowering Time

Previous studies have already performed QTL mapping for flowering time in these RILs (Lovell et al. 2013; Dittmar et al. 2014). We confirmed that the major-effect QTL identified on chromosome 4 corresponding to the gene *FRIGIDA* is recovered when using our QTL mapping workflow and phenotypes collected under our greenhouse conditions (Supplementary Figure 4a, Supplementary Table 4 in File S2). For Belm-12 x Roda-47 RILs, we confirmed that the QTL on chromosome 5 corresponding to *FLOWERING LOCUS C* is recovered with our QTL mapping workflow applied to flowering time data under Italy growth chamber conditions from Dittmar et al. (Supplementary Figure 4b, Supplementary Table 4 in File S2).

### Tests for Separate QTLs on Chromosome 5

We found support for separate QTLs underlying short stamen number and flowering time (likelihood ratio test with 1000 parametric bootstrap replicates, *P* < 0.001, Figure 4a). We were unable to reject the null hypothesis that a pleiotropic QTL underlies ovule number and either of the other two traits (*P* = 0.500 for ovule number and flowering time; *P* = 0.588 for ovule number and short stamen number; Figure 4b, 4c). The 1.5 LOD interval for the ovule number QTL is large (0-11.97 cM, Table 4) and contains the intervals for both other traits. This feature, or real pleiotropy between ovule number and stamen number or flowering time, prevents us from detecting distinct QTLs contributing to ovule number and the other traits.

**Figure 4.**
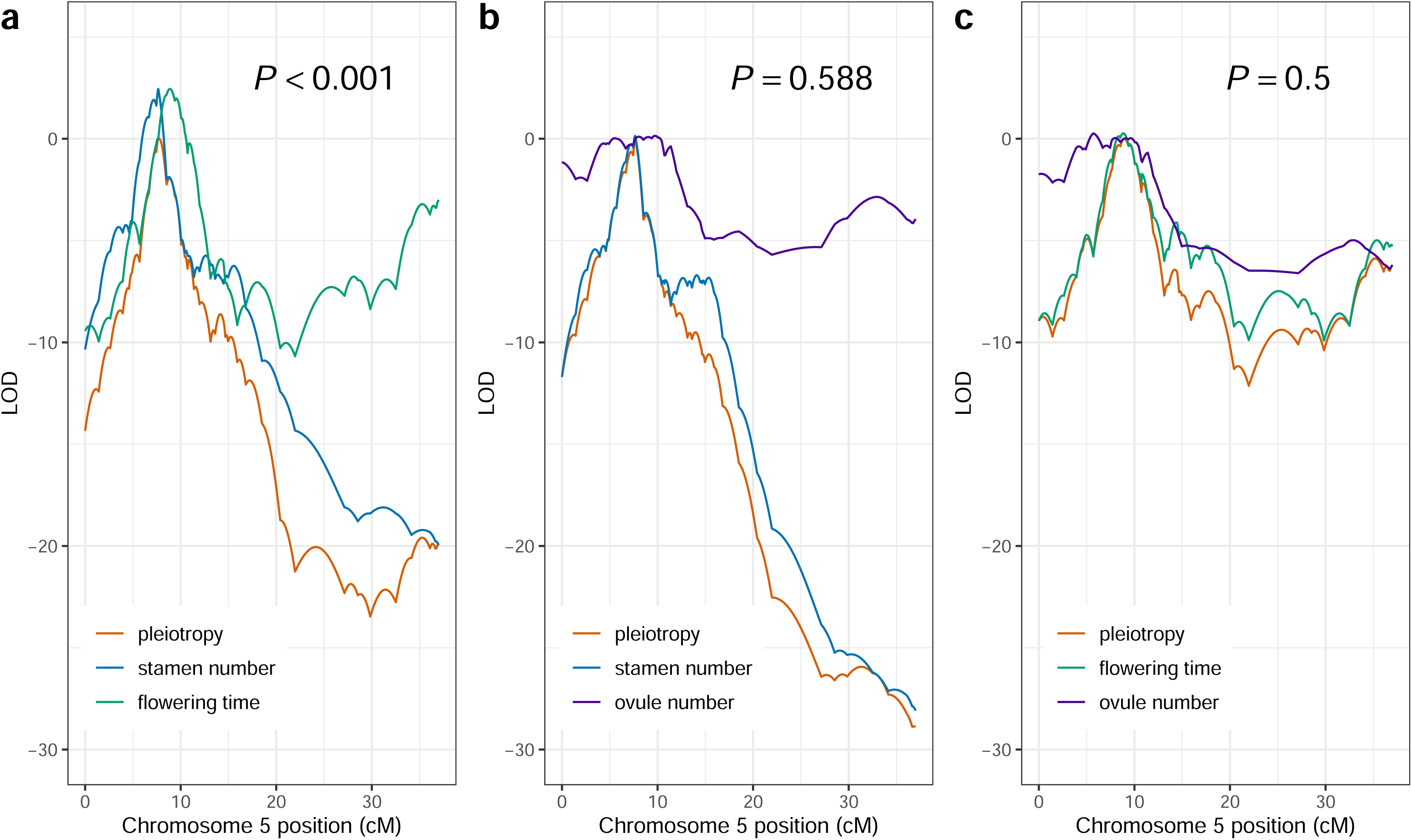
LOD plots for separate QTLs or a single pleiotropic QTL on chromosome 5 for each pair of traits in Belm-12 x Roda-47 RILs. Red-orange traces correspond to the pleiotropy LOD values for the two individual traits shown in each plot. *P-*values indicate the results of 1000 parametric bootstrap replicates, where low *P-*values are evidence of two separate QTLs underlying the traits. (a) LOD traces for short stamen number, flowering time, and a single pleiotropic locus affecting both traits. (b) LOD traces for short stamen number, ovule number, and a single pleiotropic locus affecting both traits. (c) LOD traces for flowering time, ovule number, and a single pleiotropic locus affecting both traits. Alt text: LOD plots for each pairwise combination of traits are shown, with genetic map positions 0 to 30 cM on chromosome 5 on the x-axis and LOD values from −30 to 5 on the y-axis. (a) The peaks of the stamen number and flowering time LOD traces are higher than for pleiotropy, with the stamen number, pleiotropy, and flowering time traces peaking at approximately 7, 7.5, and 9 cM respectively. *P* < 0.001. (b) Pleiotropy and stamen number traces both peak at approximately 8.5 cM, with the ovule number peak extremely broad (greater than 5 cM) and overlapping the other traces. All peaks are similar in height. *P* = 0.588. (c) Flowering time and pleiotropy traces both peak at approximately 9 cM, with the ovule number peak extremely broad (greater than 5 cM) and overlapping the other traces. All peaks are similar in height. *P* = 0.500.

### Identifying and Testing Candidate Genes

Since both mapping populations had large-effect stamen loss QTLs on chromosome 5, we sought to identify the gene underlying this QTL. We selected an interval that contains the QTL region for Belm-12 x Roda-47 but is about 0.5 Mb away from the interval for Tsu-1 x Kas-1 RILs. We prioritized the Belm-12 x Roda-47 interval due to the aforementioned effects of analysis choices on Tsu-1 x Kas-1 QTL number and position on chromosome 5 (see above section *Genetic Architecture of Short Stamen Number*), which may reflect issues in marker order and/or genomic structural divergence from Col-0. We also note that the flanking markers of the 1.5 LOD interval for the 5@3.0 cM QTL in Tsu-1 x Kas-1 RILs have increasing position in cM but decreasing position in bp (Table 3). This uncertainty about the base-pair coordinates of the 5@3.0 cM QTL led us to select candidate genes primarily in and near the Belm-12 x Roda-47 chromosome 5 QTL region.

To investigate the molecular identity of the chromosome 5 QTL, we selected 13 candidate stamen-loss genes within and near the region of chromosome 5 where stamen number QTLs were identified, from 1.5 – 3 Mb of the TAIR10 reference genome (Figure 5, Table 5). There are 482 annotated genes in this region in the TAIR10 reference genome. Twelve of our 13 chosen candidate genes are those expressed in floral tissues for which we identified mutations shared by stamen-loss parents that are expected to affect protein function, and for which T-DNA insertional mutants were available (see Methods). The thirteenth gene, *FLOWERING LOCUS C* (*FLC*), was selected as a strong candidate gene for the Belm-12 x Roda-47 flowering time QTL (Dittmar et al. 2014) for which up- and down-stream genes are known to affect stamen elongation and flower morphology (Song et al. 2022; Romera-Branchat et al. 2025). For each gene, we phenotyped short stamen number in 4-5 individuals from each line homozygous for the T-DNA insertion. Wild-type Col-0 is mostly stamen-retaining, with a mean short stamen number per plant of 1.92 that reflects occasional flowers with stamen loss. No mutant lines showed a significantly different mean short stamen number from Col-0 (Supplementary Table 2 in File S1). Additionally, all effect sizes were smaller in magnitude than the 0.4 short stamens expected for homozygotes for stamen loss alleles of QTLs detected on chromosome 5 (Table 3). Although one mutant line for MYB76 showed a marginally significant *increase* in short stamen number compared to wild-type (effect = 0.10, *P* = 0.0584), we found a non-significant loss when phenotyping this line again (effect = −0.11, *P* = 0.228), nor did we observe an increase in short stamen number in the other line we examined for this gene. Our search for genes underlying the chromosome 5 stamen loss QTL did not confirm any candidate gene that leads to stamen loss when knocked out in the Col-0 background.

**Figure 5.**
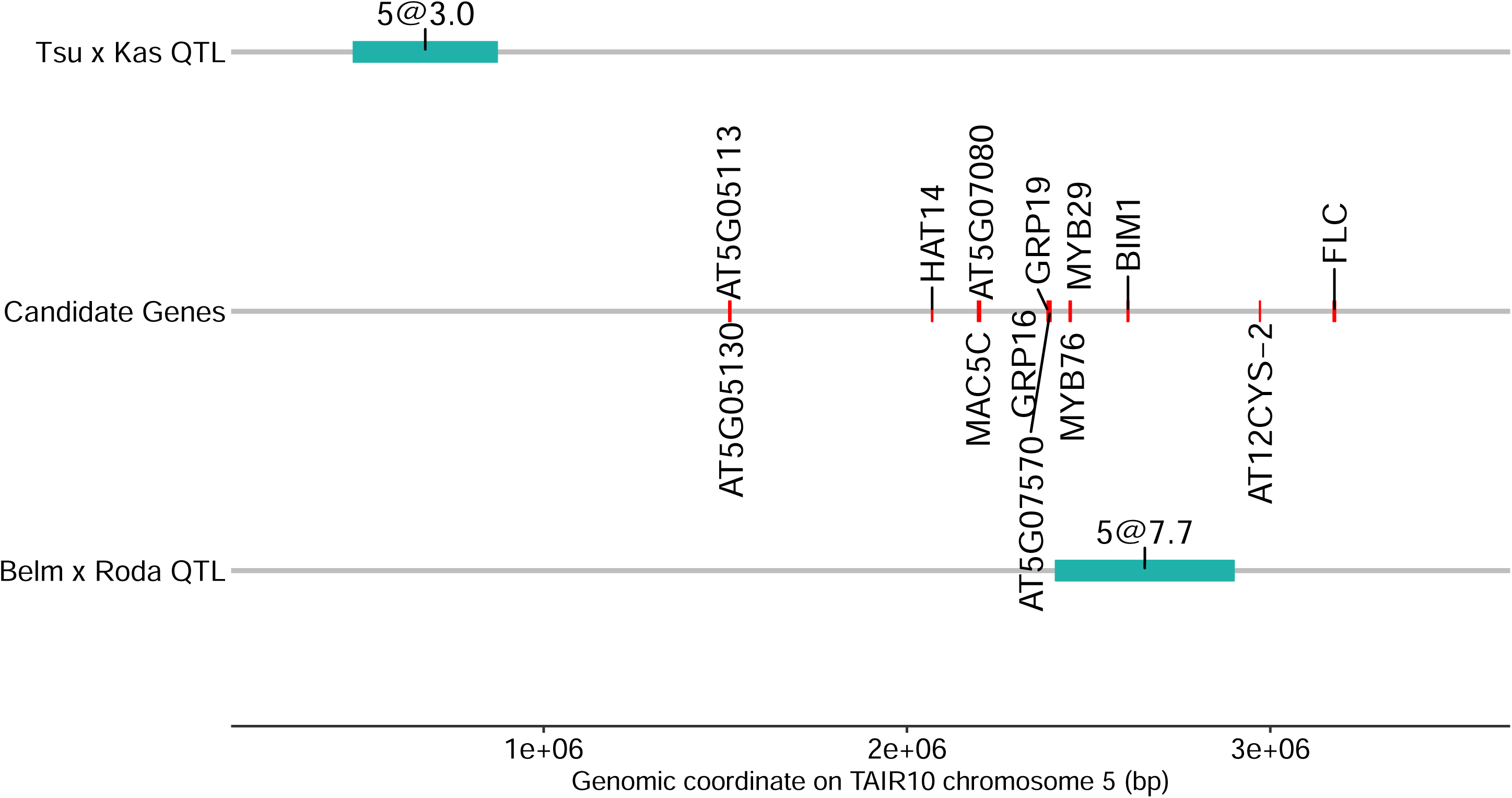
Genomic region of stamen number QTLs on chromosome 5. 1.5 LOD support interval shown in teal for Tsu-1 x Kas-1 QTL (top), candidate genes tested using TDNA insertional mutants, and 1.5-LOD support interval shown in teal for Belm x Roda QTL (bottom). Alt text: Physical genomic coordinates of chromosome 5 QTLs for both sets of RILs (Tsu-1 x Kas-1, peak at 3 cM, and Belm-12 x Roda-47, peak at 7.7 cM) are shown as teal bars about 1.5 megabases apart. Genomic positions of 14 candidate genes tested with T-DNA insertional mutants (listed in Table 5) are shown, with most of the genes falling between the two QTLs or within the Belm-12 x Roda-47 QTL.

**Table 5.** Candidate genes tested using TDNA insertional mutants. Variants column describes variants that led us to choose this candidate gene for study, for which stamen-loss parents Belm-12 and Kas-1 have a nonreference allele and stamen-retaining parents Roda-47 and Tsu-1 have a reference allele. Other rationale for selecting these genes includes expression data (Klepikova et al. 2016) and literature evidence connecting these genes or gene families to flower, stamen, or pollen development.

| Gene ID | Gene annotation | Variants | Other rationale |
|---|---|---|---|
| AT5G05113 | Hypothetical protein | Substitution leading to stop lost, 9 downstream variants, 9 upstream variants | Expressed in flower buds |
| AT5G05130 | DNA/RNA helicase protein | 16 upstream variants, 2 missense variants, 1 downstream variant | Expressed in stamens, flower buds |
| AT5G06710 | HOMEOBOX FROM ARABIDOPSIS THALIANA14 | 24 upstream variants |  |
| AT5G07060 | MOS4-associated complex subunit 5C | 3 missense variants, 8 upstream variants, 9 intronic variants | Expressed in stamens, flower buds |
| AT5G07080 | BAHD acyltransferase | 9 missense variants, 10 upstream variants, 3 intronic variants | Modulating the polyamine pathway often leads to aberrant stamen number (Kakkar and Sawhney 2002). |
| AT5G07540 | GLYCINE-RICH PROTEIN 16 | 60-bp in-frame deletion, 3 downstream variants, 4 upstream variants | Expressed in flower buds. Part of gene cluster required for normal pollen development (Lu et al. 2020). |
| AT5G07550 | GLYCINE-RICH PROTEIN 19 | 5 downstream and 2 3’ UTR variants | Expressed in stamens and flower buds. Part of gene cluster required for normal pollen development (Lu et al. 2020). |
| AT5G07570 | Glycine/proline-rich protein | 15-bp in-frame deletion, 4 upstream variants, 1 downstream variant | Expressed in flower buds. Similar to gene cluster required for normal pollen development (Lu et al. 2020). |
| AT5G07690 | MYB29 | 8 missense variants, 5 3’ UTR variants, 20 downstream variants |  |
| AT5G07700 | MYB76 | 31 upstream variants |  |
| AT5G08130 | BES1-INTERACTING MYC-LIKE1 | 60 bp deletion upstream | Part of the brassinosteroid pathway, which affects male fertility (Ye et al. 2010). |
| AT5G09570 | AT12CYS-2 | 1 intron and 1 upstream variant | Putatively regulated by stamen identity genes PI and AP3 (Mara 2008). |
| AT5G10140 | FLOWERING LOCUS C | 1 downstream, 3 intronic, and 2 upstream variants | Regulator of flowering time. Downstream (Romera-Branchat et al. 2025) and upstream (Song et al. 2022) genes affect stamen initiation and elongation. |

## Discussion

In this study, we identified shared QTL regions on chromosomes 1 and 5 that affect short stamen number in replicate mapping populations. Other stamen number QTLs were unique. Co-localization between stamen number QTLs and other trait QTLs occurred only for the correlated ovule number and flowering time traits in the Belm-12 x Roda-47 cross. These correlations were reduced in magnitude but still significant after controlling for the fraction of the RIL genome contributed by each parent. Finally, we took a novel approach to selecting candidate genes by leveraging shared variants found in the stamen-loss parents but not the stamen-retaining parents of the two RIL sets to identify 13 candidate stamen loss genes based on shared variants. However, none of these knockouts showed a significant change in short stamen number, so the molecular identity of this stamen loss QTL remains elusive.

There are several possible reasons for the lack of significant knockout effects. First, while testing candidate genes in a different genetic background (Col-0) from our RIL parents allowed us to take advantage of libraries of T-DNA insertional mutants, Col-0 may have other loci in the genome that mask the effect of mutations in the chromosome 5 stamen loss genes. The effects of mutations on phenotypes are often different between genetic backgrounds of the same species (Chandler et al. 2013); indeed, the epistatic interaction between QTLs on chromosomes 3 and 5 in Belm-12 x Roda-47 RILs is one example of this context-dependence (Royer et al. 2016). Second, changing a stamen retaining allele to a stamen loss allele may not involve a simple knockout, but rather functional changes in protein sequence or gene expression. Finally, our study design is not sufficiently replicated to detect the effect sizes expected if two or more linked loci of smaller effect underlie the chromosome 5 QTL region instead a single larger effect QTL (Flint and Mackay 2009). Verifying whether two linked QTLs are present and determining their molecular identity using fine-mapping and transgenics in the genetic backgrounds of the RIL parental lines are critical next steps to uncover the genes underlying these small-effect stamen number loci. Indeed, the chromosome 5 QTL region contains multiple genes in the *GLYCINE RICH PROTEIN* (*GRP*) family that are expressed in flower buds and form part of a gene cluster required for normal pollen development (Lu et al. 2020, Table 5). Multi-gene functional studies, such as higher-order mutant analysis or gene family knockouts (e.g. Anfang et al. 2026), will be key to understanding whether such clustered gene families within QTLs contribute redundantly to traits like stamen loss.

Our study supports a cautious approach to interpreting trait correlations in mapping populations as evidence of linkage or pleiotropy. Individuals in a mapping population can display substantial variation in the fraction of the genome contributed by each parent (Supplementary Figure 2 in File S2, Frisch and Melchinger 2007). If the Belm-12 and Roda-47 parents have contrasting phenotypes for multiple traits, and the individuals in the mapping population inherit variable amounts of the Roda-47 genome, then we may expect RILs that inherited a large amount of the Roda-47 genome to display multiple phenotypes resembling Roda-47, even without linked or pleiotropic loci underlying variation in these traits. Indeed, controlling for this variation in parental genome contribution reduced some of the trait correlations in Belm-12 x Roda-47 RILs (Table 2). Correlations in RILs therefore indicate correlated divergence between parents, which *may* involve linked or pleiotropic loci but are not necessarily informative about the genetic mechanisms of trait correlations in nature.

The contrasting genetic architectures of flowering time in the two sets of RILs likely drives cross-specific correlations between short stamen number and flowering time. The LOD peaks for the chromosome 5 QTLs affecting flowering time and stamen number in Belm-12 x Roda-47 are less than 1 cM apart, suggesting tight linkage or pleiotropy in this cross (Table 3, Supplementary Table 4 in File S2). Knockout mutants of *flc*, the putative molecular basis of the flowering time QTL (Dittmar et al. 2014), showed no change in short stamen number (Supplementary Table 2 in File S1, and *qtl2pleio* analyses strongly support separate QTLs for these traits (Figure 5a). We therefore propose that physical linkage between these QTLs, not pleiotropy, contributes to the correlation between short stamen number and flowering time in Belm-12 x Roda-47 RILs.

In natural populations, linkage or pleiotropy with flowering time genes may lead to clines in stamen and ovule number as a correlated response to selection on flowering time. The repressor of flowering *FLC* is a strong candidate gene underlying flowering time variation in Belm-12 x Roda-47 RILs (Dittmar et al. 2014). In Tsu-1 x Kas-1 RILs, *FRIGIDA* (*FRI*) underlies a flowering time QTL at 4 cM on chromosome 4 (Lovell et al. 2013) which is not closely linked to any detected stamen loss QTL. *FLC* and *FRI* epistatically interact to produce a latitudinal cline in *A. thaliana* flowering time (Caicedo et al. 2004). Flowering time variation is thus driven both by loci like *FRI* that are unlinked to stamen number QTL, and by loci like *FLC* that are linked to the repeatable chromosome 5 stamen number QTL uncovered in both crosses.

Overlapping QTLs may also drive the negative correlation between ovule number and stamen number observed in Belm-12 x Roda-47 but not Tsu-1 x Kas-1. We identified an ovule number QTL on chromosome 5 that was unique to the Belm-12 x Roda-47 RILs and overlaps the shared stamen number QTL region on chromosome 5 (Table 3, Table 4). Individuals carrying a Roda-47 allele at these loci would have more short stamens and fewer ovules than individuals carrying a Belm-12 allele. If stamen loss alleles were commonly linked to flowering time and ovule number QTLs, such architecture would offer a potential mechanism for correlations between flowering time, ovule number, and stamen number in natural accessions (Royer et al. 2016). Physical linkage of stamen- and ovule-number QTLs with *FLC* offers a potential mechanism for observed latitudinal clines in these traits (Royer et al. 2016), but note Yuan and Kessler (2019) reported a lack of a latitudinal cline in ovule number. *qtl2pleio* analysis did not support separate loci affecting ovule number and short stamen number, nor ovule number and flowering time. Further fine mapping and functional validation are required to understand the roles of linkage and pleiotropy in ovule number variation.

Just as linkage between flowering time, ovule number, and short stamen number QTLs on chromosome 5 in Belm-12 x Roda-47 RILs suggests a mechanism for correlations in natural accessions, the lack of large-effect overlapping QTLs for these traits in Tsu-1 x Kas-1 allows those traits to evolve independently. Similarly, additional QTLs on chromosomes 3 and 5 that do not overlap with any stamen number QTLs found in this study were identified in a genome-wide association study for short stamen number across an elevational gradient (Buysse et al. 2025). Our QTL analysis indicates that the genetic architecture of stamen and ovule number can involve multiple different loci across the genome, with some QTLs unique to a specific cross. Consistent with other QTL studies of multiple mapping populations (Dilda and Mackay 2002; Symonds et al. 2005; Kumar et al. 2007), the genetic architectures of the studied traits are specific to certain crosses, which can in turn influence patterns of physical linkage between QTLs affecting different traits.

## Data availability

Parental sequencing reads are available on the Sequence Read Archive under BioProject PRJNA1468757. Code and phenotype data area are available on GitHub (https://github.com/plunkert/Stamen-loss-2025).

## Author contributions

MLP collected and analyzed data and wrote the original draft of the manuscript. OIS performed initial QTL analyses. MLP, EW, and SGP genotyped and phenotyped TDNA insertional mutants. JC conceived of the study, supervised the project, and secured funding. All authors read and approved the final manuscript.

## Acknowledgements

The Illumina sequencing data were acquired using instrumentation in the Genomics Core Facility (RRID:SCR_012406), supported by Michigan State University’s Office of Research & Innovation. We thank Anne Royer for phenotyping Tsu-1 x Kas-1 RILs. We thank John Lovell, Tom Juenger, and members of the Lowry and Conner labs for comments on the manuscript, and Jimmy Bingman for advice on the role of parental genome contribution in trait correlations. We thank Acer VanWallendael for advice on QTL analysis and Fred Boehm for advice on tests of pleiotropy.

