## Supplementary File 2 for "Shared QTLs underlie loss of a conserved trait in replicate mapping populations of *Arabidopsis thaliana*"

Supporting Information for Plunkert et al., Shared QTLs underlie loss of a conserved trait

**Supplementary Table 1.** Primers used in this study. Primers are listed by an arbitrary ID and their sequences are indicated from 5’ to 3’. The purpose of each primer is indicated as LP (primer binds left of TDNA insertion site) or RP (primer binds right of TDNA insertion site). The LB primers bind the left border of the TDNA insertion. For each transgene, a band produced by PCR with LP and RP primers indicates that the transgene is absent, and a band produced by the LB and LP primer indicates that the transgene is present.

**Supplementary Table 2**. Effects of T-DNA insertional mutations on short stamen number. All rows report the effect of the indicated genotype (as a line shown to be homozygous for T-DNA insertion in the indicated gene) on short stamen number, compared to wild-type Col-0 grown in the same tray.

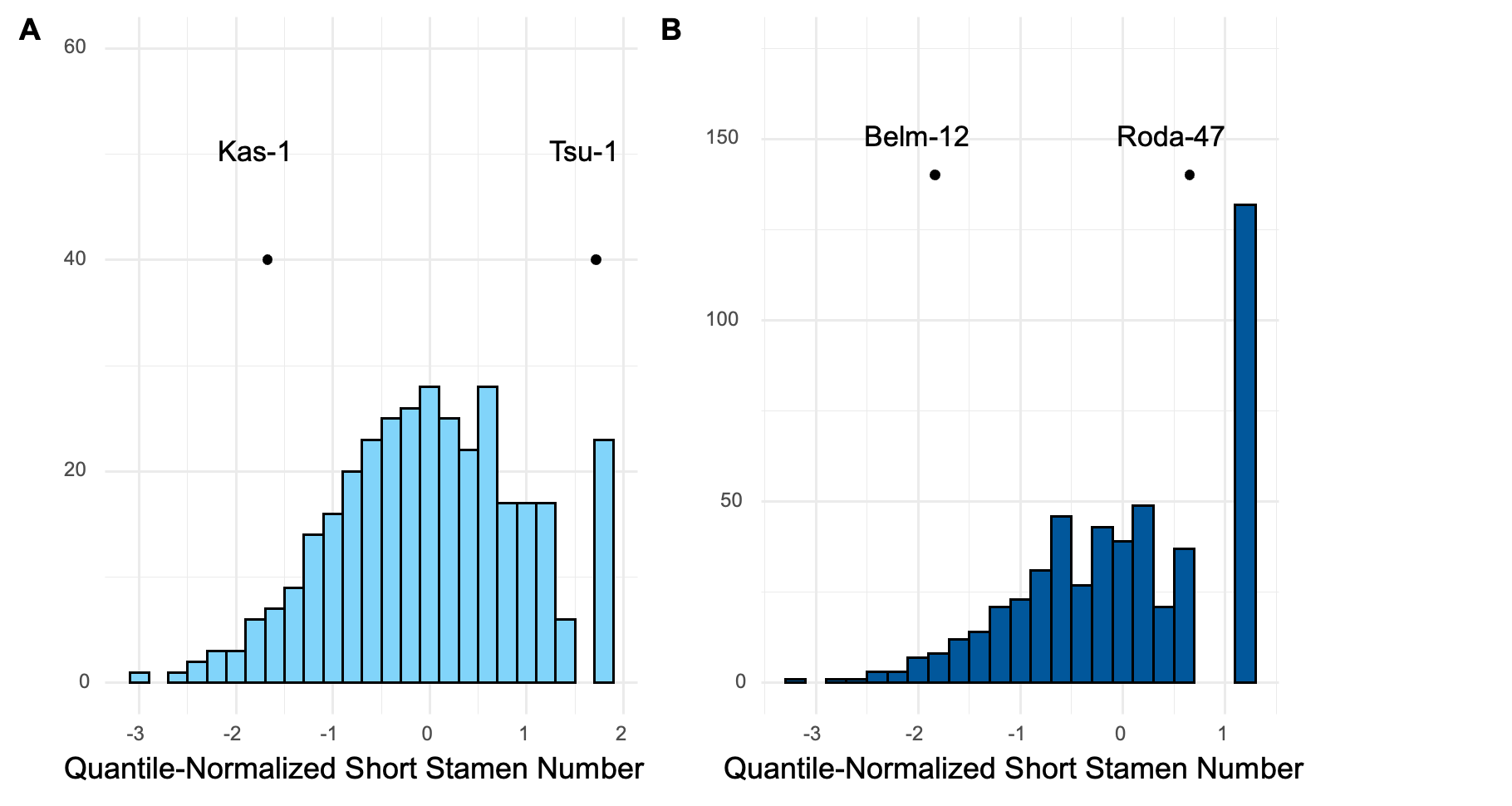

**Supplementary Figure 1.** Distribution of quantile-normalized short stamen number for Tsu-1 x Kas-1 RILs (A) and Belm-12 x Roda-47 RILs (B). Points indicate mean phenotypes of parental accessions following quantile normalization.

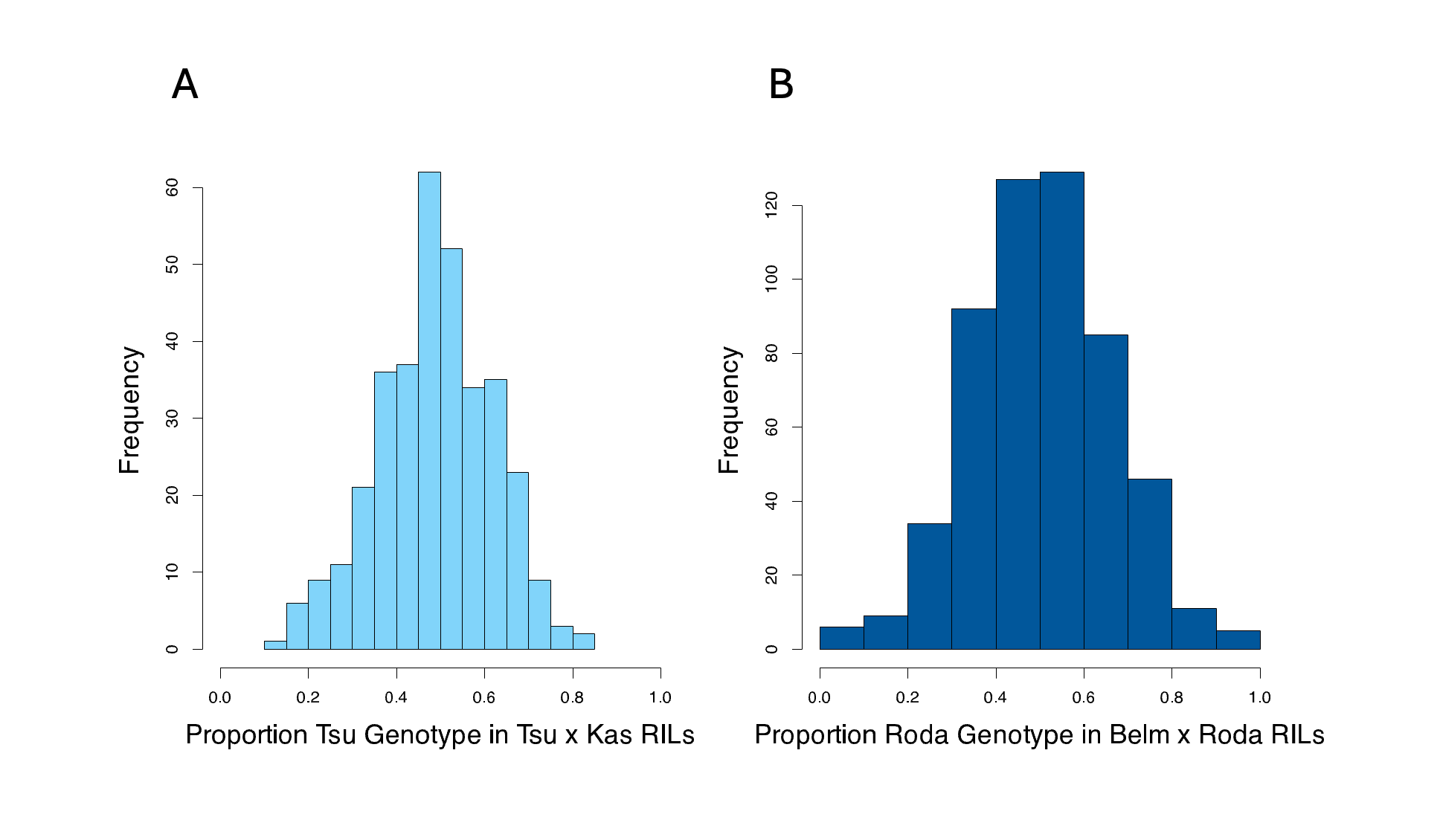

**Supplementary Figure 2.** Distribution of estimated proportion of RIL genome contributed by Tsu-1 parent in 341 Tsu-1 x Kas-1 RILs (A) and by Roda-47 parent in 544 Belm-12 x Roda-47 RILs (B).

**Supplementary Table 3.** Quantile-normalized short stamen number QTL from both mapping populations. The additive effect on quantile-normalized short stamen number for one allele from the stamen-retaining parent is given in a, %var is the percent of RIL variation explained by that QTL, and %div is percent of parental divergence explained, as 100 * 2a / (stamen retaining parent mean short stamen number – stamen loss parent mean short stamen number).

Tsu-1 x Kas-1 RILs

| QTL | Chr | Position (cM) | 1.5 LOD Interval (cM) | 1.5 LOD Interval (Mb) | LOD | a | %var | %div |
| --- | --- | --- | --- | --- | --- | --- | --- | --- |
| 1@33.6 | 1 | 33.61 | 30.15 – 65 | 9.34 – 19.58 | 3.71 | 0.15 | 1.97 | 8.91 |
| 1@80.0 | 1 | 80 | 77 – 91 | 22.68 – 25.50 | 8.17 | 0.24 | 4.47 | 14.36 |
| 1@108.0 | 1 | 108 | 107 – 108.94 | 29.79 – 28.67 | 23.03 | 0.41 | 14 | 24.41 |
| 3@86.0 | 3 | 85.98 | 76 – 89.5 | 20.20 – 21.63 | 9.58 | -0.23 | 5.3 | -13.61 |
| 4@56.1 | 4 | 56.09 | 52 – 66 | 9.32 – 14.39 | 4.74 | 0.16 | 2.54 | 9.72 |
| 5@3.0 | 5 | 3 | 2.01 – 3.80 | 0.87 – 0.48 | 34.26 | 0.49 | 22.59 | 29.02 |
| Interaction 1@108.0:5@3.0 | | | | | 1.45 | -0.1 | 0.76 | -5.61 |

Belm-12 x Roda-47 RILs

| QTL | Chr | Position (cM) | 1.5 LOD Interval (cM) | 1.5 LOD Interval (Mb) | LOD | a | %var | %div |
| --- | --- | --- | --- | --- | --- | --- | --- | --- |
| 1@74.4 | 1 | 74.44 | 67 – 80.89 | 25.03 – 29.08 | 4.23 | 0.14 | 2.31 | 11.4 |
| 3@19.7 | 3 | 19.71 | 18.03 – 22 | 7.91 – 8.50 | 12.67 | 0.23 | 7.2 | 18.49 |
| 5@7.7 | 5 | 7.65 | 6.68 – 8.53 | 2.41 – 3.27 | 48.18 | 0.54 | 32.27 | 42.99 |
| Interaction 3@19.7:5@7.7 | | | | | 4.05 | -0.14 | 2.21 | -11.4 |

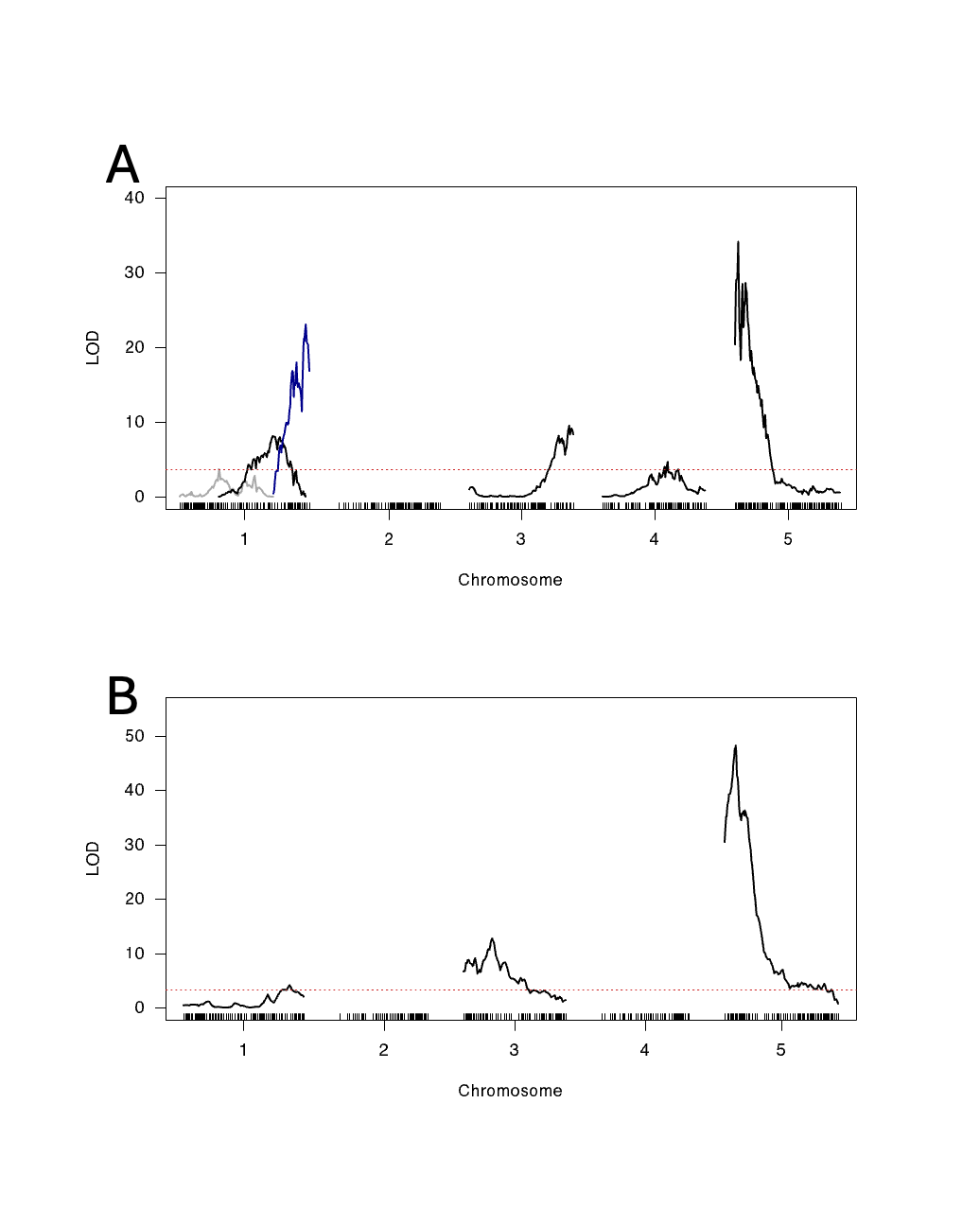

**Supplementary Figure 3**. LOD plots showing multi-QTL mapping of short stamen number following quantile normalization for Tsu-1 x Kas-1 (A) and Belm-12 x Roda-47 (B) mapping populations. Red dashed lines indicate thresholds from scantwo permutations at significance level of 0.01. Each trace indicates the LOD profile for a separate QTL, with colors used to distinguish QTL traces on the same chromosome (see Supplementary Table 3).

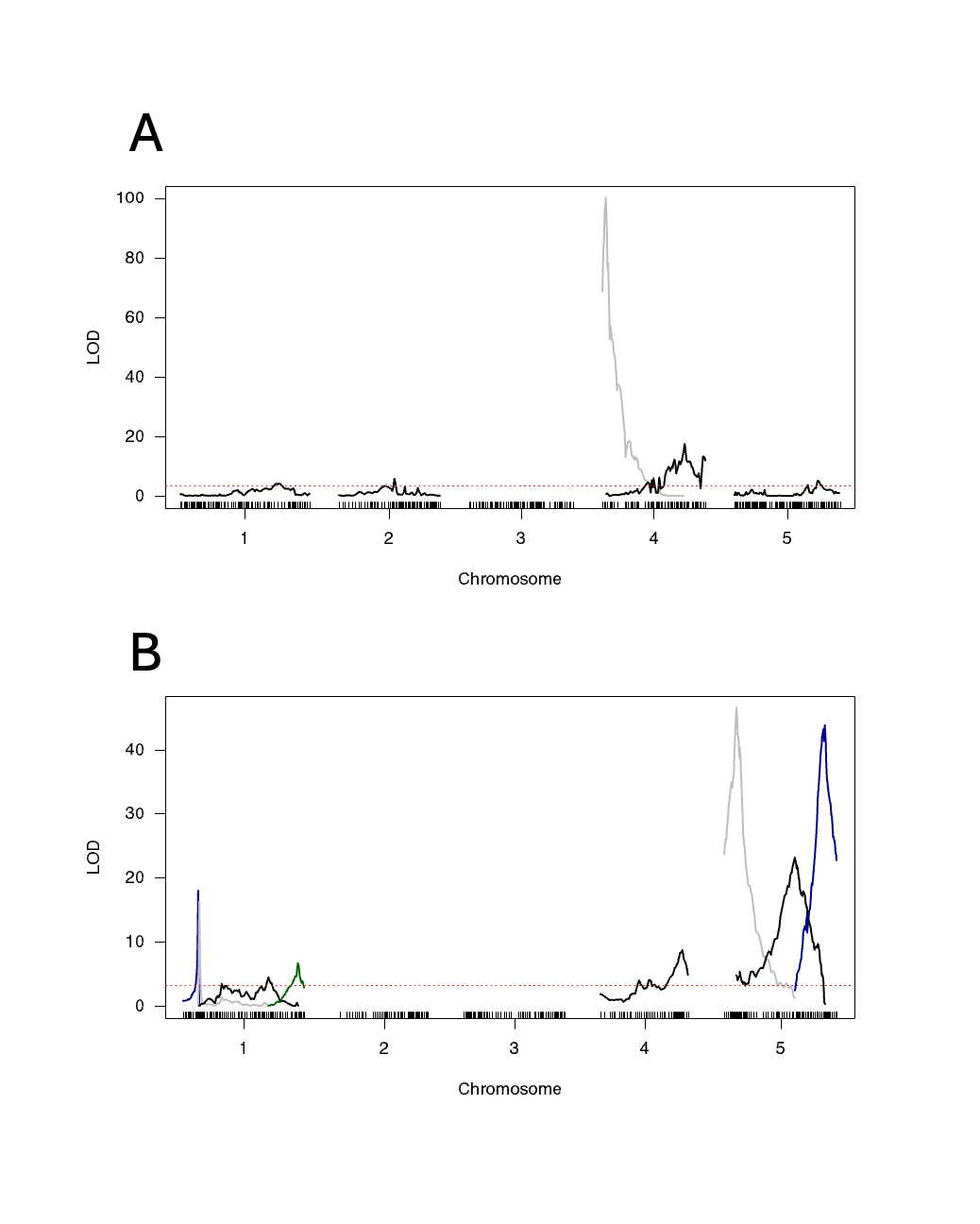

**Supplementary Figure 4**. LOD plots showing multi-QTL mapping of flowering time for Tsu-1 x Kas-1 (A) and Belm-12 x Roda-47 (B) mapping populations. Red dashed lines indicate thresholds from scantwo permutations at significance level of 0.01. Each trace indicates the LOD profile for a separate QTL, with colors used to distinguish QTL traces on the same chromosome.

**Supplementary Table 4**. Flowering time QTLs from both mapping populations. The additive effect on flowering time for one allele from the stamen-retaining parent is given in a, %var is the percent of RIL variation explained by that QTL, and %div is percent of parental divergence explained, as 100 * 2a / (stamen retaining parent mean flowering time – stamen loss parent mean flowering time).

Tsu-1 x Kas-1 RILs

| QTL | Chr | Position (cM) | 1.5 LOD Interval (cM) | 1.5 LOD Interval (Mb) | LOD | a | %var | %div |
| --- | --- | --- | --- | --- | --- | --- | --- | --- |
| 1@85.0 | 1 | 85.00 | 78.47 – 89 | 23.42 – 25.59 | 4.13 | 1.90 | 1.42 | -17.05 |
| 2@47.7 | 2 | 47.70 | 47 – 49 | 9.37 – 9.26 | 5.68 | -2.11 | 1.97 | 18.91 |
| 4@3.0 | 4 | 3.00 | 2.79 – 4 | 0.21 – 0.07 | 100.10 | -12.97 | 71.31 | 116.35 |
| 4@70.7 | 4 | 70.75 | 70.16 – 72 | 14.77 – 15.29 | 17.42 | -3.22 | 6.58 | 28.90 |
| 5@72.0 | 5 | 72.00 | 62 – 73.75 | 18.32 – 22.64 | 5.22 | 2.08 | 1.91 | -18.64 |
| Interaction 4@3.0:4@70.7 | | | | | 7.10 | 2.44 | 2.49 | -21.89 |

| QTL | Chr | Position (cM) | 1.5 LOD Interval (cM) | 1.5 LOD Interval (Mb) | LOD | a | %var | %div |
| --- | --- | --- | --- | --- | --- | --- | --- | --- |
| 1@10.3 | 1 | 10.31 | 10 – 11 | 5.01 – 5.59 | 17.93 | 3.47 | 4.85 | 35.77 |
| 1@11.0 | 1 | 11.03 | 10.31 – 11.50 | 5.31 – 5.89 | 16.32 | -3 | 4.38 | -30.87 |
| 1@58.8 | 1 | 58.82 | 26 – 62 | 10.52 – 23. 86 | 4.5 | 0.44 | 1.15 | 4.58 |
| 1@79.6 | 1 | 79.56 | 78.55 – 80.89 | 28.22 – 29.01 | 6.73 | 0.54 | 1.73 | 5.54 |
| 4@56.6 | 4 | 56.59 | 53.54 – 59 | 14.87 – 17.80 | 8.69 | 0.47 | 2.26 | 4.81 |
| 5@8.5 | 5 | 8.53 | 8 – 9 | 2.99 – 3.57 | 46.54 | 1.46 | 14.37 | 15.01 |
| 5@49.0 | 5 | 49 | 47 – 50.96 | 15.46 – 17.50 | 23.18 | 0.91 | 6.43 | 9.37 |
| 5@70.0 | 5 | 70 | 68 – 70.60 | 23.01 – 24.17 | 43.8 | 1.51 | 13.35 | 15.56 |
| Interaction 1@10.3:1@11.0 | | | | | 14.87 | 3.75 | 3.97 | 38.64 |
| Interaction 4@56.6:5@49.0 | | | | | 4.12 | -0.4 | 1.05 | -4.08 |
| Interaction 5@8.5:5@70.0 | | | | | 5.13 | -0.43 | 1.31 | -4.43 |

Belm-12 x Roda-47 RILs

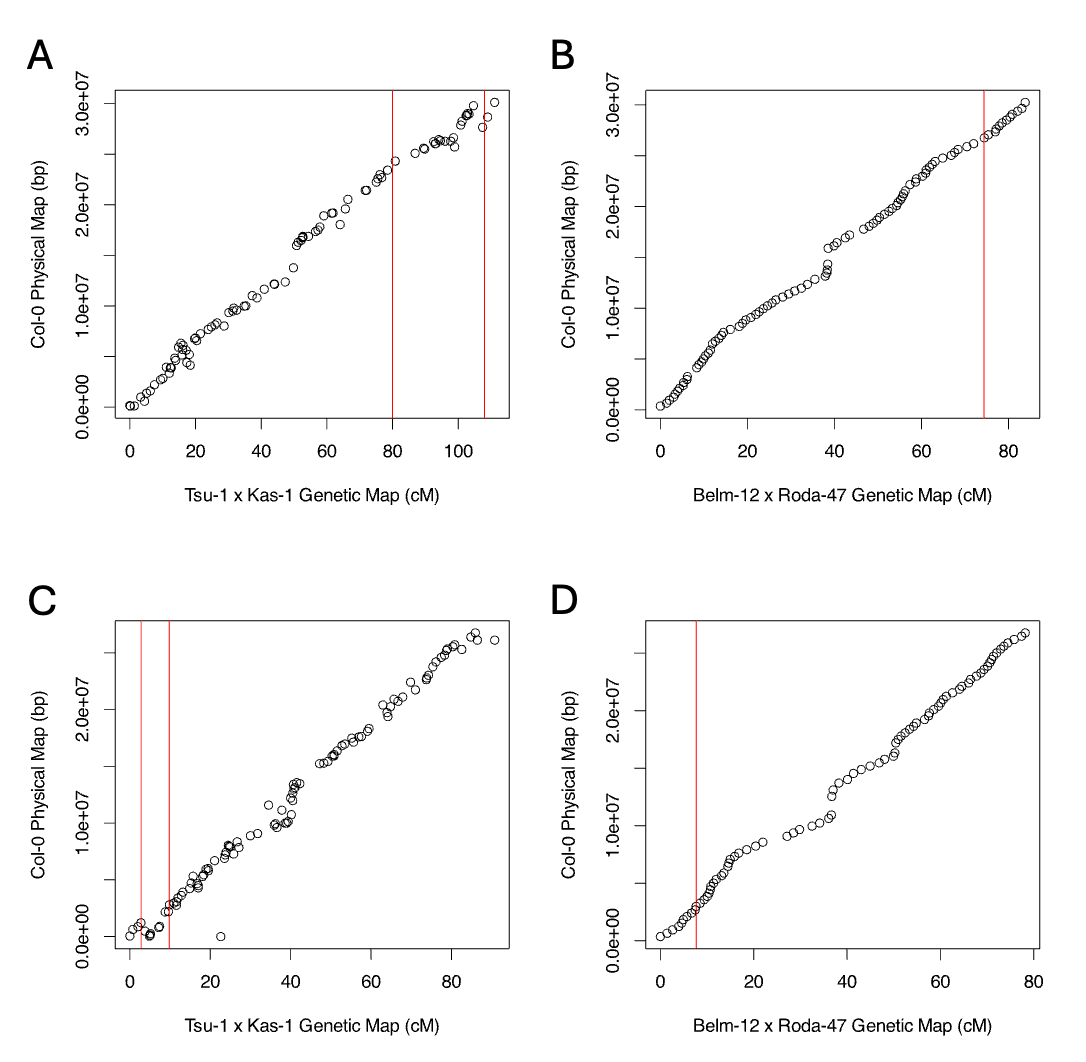

**Supplementary Figure 5.** Scatter plots showing position of each marker in the TAIR10 reference genome on the y-axis, and the marker position in the genetic maps for chromosome 1 in Tsu-1 x Kas-1 (A), chromosome 1 in Belm-12 x Roda-47 (B), chromosome 5 in Tsu-1 x Kas-1 (C), and chromosome 5 in Belm-12 x Roda-47 (D). The peaks of each short stamen number QTL detected using stepwise mapping are indicated with red lines. In (C), the positions at 2.8 and 9.8 cM (corresponding to an initial stepwise analysis before dropping the QTL at 9.8 cM) are shown.
